# Toffee – a highly efficient, lossless file format for DIA-MS

**DOI:** 10.1101/628933

**Authors:** Brett Tully

## Abstract

The closed nature of vendor file formats in mass spectrometry is a significant barrier to progress in developing robust bioinformatics software. In response, the community has developed the open mzML format, implemented in XML and based on controlled vocabularies. Widely adopted, mzML is an important step forward; however, it suffers from two challenges that are particularly apparent as the field moves to high-throughput proteomics: large increase in file size, and a largely sequential I/O access pattern.

Described here is ‘toffee’, an open, random I/O format backed by HDF5, with lossless compression that gives file sizes similar to the original vendor format and can be reconverted back to mzML without penalty. It is shown that mzML and toffee are equivalent when processing data using OpenSWATH algorithms, in additional to novel applications that are enabled by new data access patterns. For instance, a peptide-centric deep-learning pipeline for peptide identification is proposed.

Documentation and examples are available at https://toffee.readthedocs.io, and all code is MIT licensed at https://bitbucket.org/cmriprocan/toffee.

## Introduction

Biobank-scale data independent acquisition mass spectrometry (DIA-MS) poses a significant set of analytics challenges. In particular, challenges such as scale, automation, data management, and reproducibility are exacerbated by proprietary data formats that pose a barrier to efficient analysis and re-analysis of data over time.

In response, the community has developed many new and open data formats. Some, such as mzML^1^, and mz5^2^, aim to be archival formats that faithfully adopt the HUPO PSI guidelines. In doing so, these formats adhere to recording data in vectors of mass over charge (m/z) and intensity pairs, with one vector for each scan of the mass spectrometer (at a recorded retention time). Typically, software will access the data in these files one scan at a time, loading all of the data for a scan before moving to the next scan. Performance gains are made by indexing the scans so that the file need not be read from the beginning each time. Regardless, algorithms are limited to taking slices of the data along the retention time (or spectrum) axis only. In contrast, other attempts such as mzDB^3^, mzRTree^4^, and mzTree^5^, focus on random I/O access through the use of an RTree^6^ data structure. This allows the data to be accessed along both the m/z (or chromatogram) axis as well as the retention time axis, at the cost of file size or mass accuracy.

The motivation of the current work was to combine the benefits of a variety of these approaches, selecting those specifically geared towards minimizing the challenges of analysis and re-analysis of biobank-scale DIA-MS data. Generally, these benefits can be grouped into two categories: file size, and data access through open protocols. Here, I outline toffee, a new file format and software packages for interacting with the raw data efficiently.

## Toffee

Born out of The ACRF International Centre for the Proteome of Human Cancer (ProCan), toffee has been primarily developed for data generated on Sciex instruments, collecting data in DIA-MS mode. ProCan aims to collect data from up to 150,000 mass spectrometer injections by 2025, thus the challenges of current file formats outlined above are acute.

### Open protocol file size

Biobank-scale proteomics facilities may run upwards of 100,000 SWATH-MS runs; operating in a manner typical to ProCan results in Sciex `wiff` files 1-2 GB per injection that unpack to 10-20 GB when converted to `mzML` and lead to petabytes of data that must be kept available for long-term re-analysis. Furthermore, this increase in file size adds significant analytics time as software becomes largely IO-bound.

### Random access

Indexed mzML substantially improves randomly accessing single scan data (at constant retention time), yet algorithms often require slices along the m/z axis and thus necessitate expensive iteration of the full mzML file or high-memory compute infrastructure to keep data in RAM.

### Testability

A key challenge to improving downstream software is the slow iterative cycle imposed by storing experimental data in mzML. Building reliable and robust algorithms requires a strong testing framework of both unit and regression tests and a test harness that encourages developers to use it. The IO-bound nature of mzML files raises artificial barriers to test adoption.

Toffee addresses each of these challenges through a lossless compression mechanism, the open HDF5^7^ storage protocol, and a `boost::rtree` run time implementation. Importantly, it does not aim to be a long-term *archival* format and takes the view that this is the role of the original vendor file. Instead, toffee fills the role of efficient raw data access through open protocols, and although it does not currently implement the PSI ontology, attributes and metadata can be added trivially should the user wish to do so. While vendor lock-in for archiving is a disadvantage, the highly regulated clinical environment in which vendors operate suggests that there *should* always be a way to access the data within these files if necessary.

### Lossless compression

The following explanation of how toffee compression is achieved reflects its heritage of development on Sciex TOF data. However, the fundamental approach of converting m/z to an integer index space should be applicable to data collected on both time-of-flight, and Orbitrap mass analyzers. In making toffee open-source, contributions on its extension by others more familiar with different instruments are highly encouraged. In particular, I anticipate it is extremely well suited to include ion mobility for efficient analysis of timsTOF DIA-MS data.

Toffee’s lossless compression is achieved by understanding the physics of a time-of-flight (TOF) mass analyzer. Here, a charged ion is accelerated through an electric potential; by applying Maxwell’s equations on the work done on the ion and Newton’s laws of motion, you can say that the m/z of the ion is related to its time of arrival at the sensor through the following relationship:

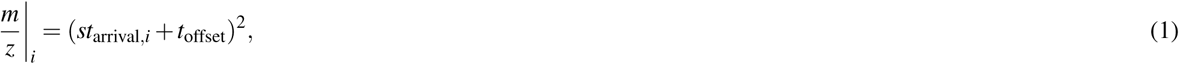

where the arrival time is a multiple of the sensor sampling rate (Δ*t*) plus an offset (*t*_0_) such that

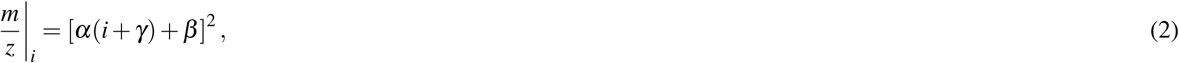

where *α* = *s*Δ*t, β* = *t*_offset_ + *st*_0_ − *αγ*, and *γ* are the Intrinsic Mass Properties (IMS) of the injection^8^. For a full derivation see the Methods section.

Pragmatically, given the data of an mzML file, toffee is able to iteratively calculate the IMS properties; ideally, this data would be exposed directly from the vendor format but this cannot always be relied upon. Interestingly, in order to ensure lossless compression, the IMS parameters must be calculated and stored *for each scan* rather than for a given injection, or even scan type (i.e. MS1 or MS2). The implication here is that the mass spectrometer performs in-line calibration that is additional to the manual calibration completed as part of routine lab operation (see Supplementary Material *calculate_ims.ipynb* for more information).

From this m/z transfer function, it is possible to convert m/z values to integer m/z indices (*i*), while retention time is calculated from the scan index and the instrument cycle time. Thus, all raw data in the file can be represented as a vector of integer triplets: m/z index, retention time index, and intensity. These triplets fall onto a Cartesian grid, and thus can be stored as a compressed row storage sparse matrix^9^. This sparse matrix can be efficiently saved as three data sets in the toffee file with zlib compression provided natively by HDF5. Further details can be seen in the Methods section.

The Toffee file format is accompanied by C++ and python libraries for creating and accessing the file format. All code is MIT licensed at https://bitbucket.org/cmriprocan/toffee and documentation and examples are available at https://toffee.readthedocs.io.

## Results

### File size comparisons

File size is crucial to manage long-term data storage and retrieval costs, in addition to minimizing the hardware that is required during any computational analysis of the raw data. Using three public data sets covering both TripleTOF5600 (Swath Gold Standard^10^ and TRIC manual validation set, only the y- and b-ions are included in the analysis,^11^) and TripleTOF6600 (ProCan90, including only the first injection from each mass spectrometer,^12^) the raw vendor files were converted to mzML using ‘msconvert’^13,14^, in both profile and vendor peak-picking centroid mode, each with and without ‘msnumpress’^15^, as well as the ‘sciex/wiffconverter’ Docker image^16^ in both profile and centroid modes. Toffee files were then produced from the msconvert and Sciex Docker profile mzML files, and the toffee file back to mzML. Figure 1 shows that the largest mzML files are produced by ‘msconvert’ with no ‘msnumpress’ compression, and the smallest are either the lossy centroided and compressed mzML files, or the lossless profile toffee files, both of which compare in size to the vendor format. Finally, for reference, a small subset of files were converted to mz5.

**Figure 1.**
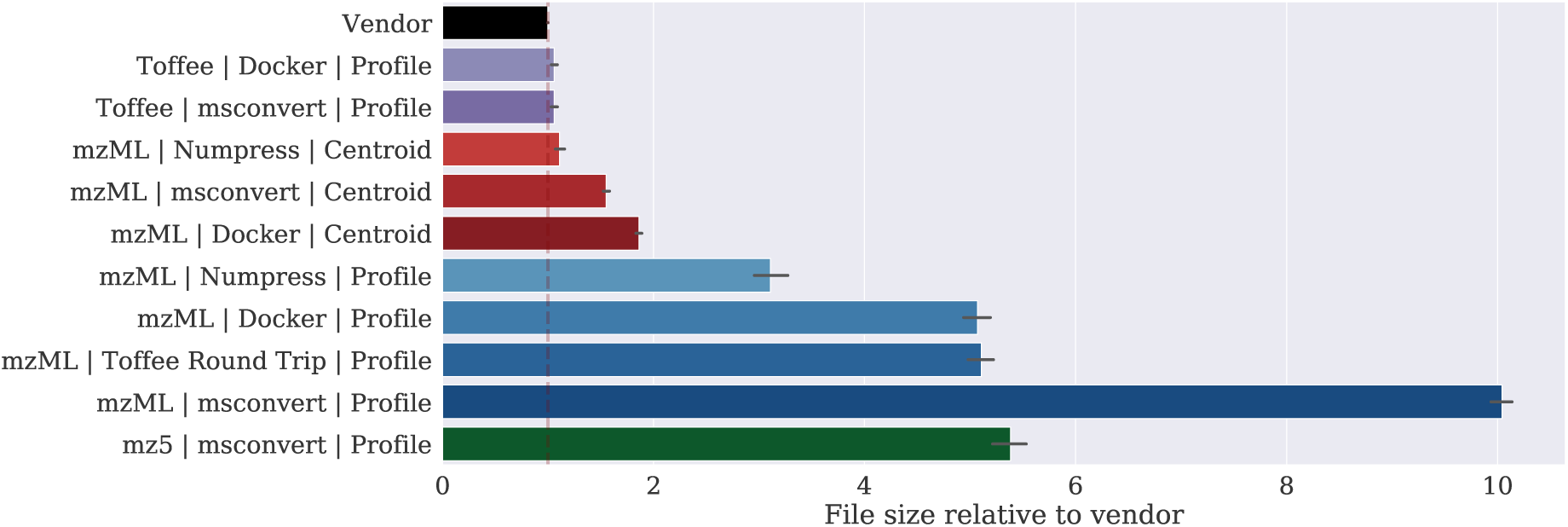
Comparison of several different file conversion methods, including: profile and centroided mzML created from ‘msconvert’ and the ‘Sciex Docker image’, mz5 files created using ‘msconvert’ and toffee files created using the method described in this work. Clearly, profile mzML files require the most storage although this can be improved based on the method of generation and if ‘numpress’ compression is used. Centroiding the data results in much smaller files, and toffee files are of equivalent size to the original vendor format. See the Methods section for a full description of all conversion methods.

### Raw data access

Enabling random, rather than sequential, data access is beneficial on multiple fronts. In peptide-centric approaches such as OpenSWATH, it is a significant algorithmic advantage to enable accessing data by slicing through the m/z axis. As Table 1 shows, compared with an indexed mzML file, toffee is around 4 times more efficient for spectrum-centric and two orders of magnitude more efficient for peptide-centric data access. This analysis was conducted on a laptop with a single thread and under minimal RAM usage (<5 GB). It is worth noting that these types of timings are somewhat artificial. The most suitable method of loading and caching raw data will be highly dependent on the application, algorithm, and its implementation – performance is always best achieved through profiling the actual code being optimized. Furthermore, there is no consideration given here to threaded access, or high memory environments that would allow all data to be held in RAM. Full details of this comparison can be seen in Supplementary Material *random-access-timing.ipynb*.

**Table 1.** Accessing data from toffee files is ∼4 times more efficient for spectrum-centric access, and ∼100 times more efficient for peptide-centric access. MS2 window “050” was selected as an indicative medium/high load window. The setup for reading mzML requires a loop through the file to extract the index and match it to its relevant MS1 and MS2 windows, while toffee must load its relevant data classes.

| Process | Time mzML (s) | Time toffee (s) |
| --- | --- | --- |
| Setup MS1 & MS2 data | 151 | **11.42** |
| Extract spectrum data from MS1 & MS2-050 | 3.99 | **0.66** |
| Extract spectrum data from MS2-050 | 1.61 | **0.41** |
| Extract all chromatogram data for MS2-050 peptides | 168 | **1.79** |

### Replacing mzML for analysis

The Toffee file format is accompanied by a wrapper around OpenSWATH^10^ that enables SWATH-MS data to be analyzed with standard algorithms and the scores to be used in False Discovery Rate (FDR) calculations of PyProphet^17^. The MIT-licensed OpenMSToffee^18^ serves two purposes: to demonstrate that toffee does not introduce any artifacts to the OpenSWATH pipeline converting raw data to a quantified list of peptides, and as an exemplar of how peptide-centric data extraction can be achieved using the C++ toffee library.

Using OpenMSToffee I have conducted a thorough investigation into OpenSWATH with a variety of mzML conversions, and toffee itself. In order to assess the quality of the data in each file conversion method, they are input into the computational pipeline as described in the Methods section (all analysis code is included in the ‘openms-toffee-paper’ repository^19^ and is executable on MacOS or Linux). One of the reasons for selecting the SGS and TRIC data sets for this analysis, is their inclusion of a collection of manually validated peptide query matches (PQMs). Using this information, the results of the file format pipelines are assessed by categorizing those results with a peptide retention time within 15 seconds of the manually validated peak as a *true positive*, those not within this threshold as a *false positive*, and those peaks in the manual validation, but not found using this file format as *false negatives*. One can see from Figure 2 that each of the profile files (mzML and toffee) performs equivalently, while centroided mzML files have a larger number of false positives particularly with the TRIC data (see Figure 2C). This is further demonstrated in Figure 2D by the increase in missing values that are seen with centroided mzML files when analyzing the ProCan90 data set.

**Figure 2.**
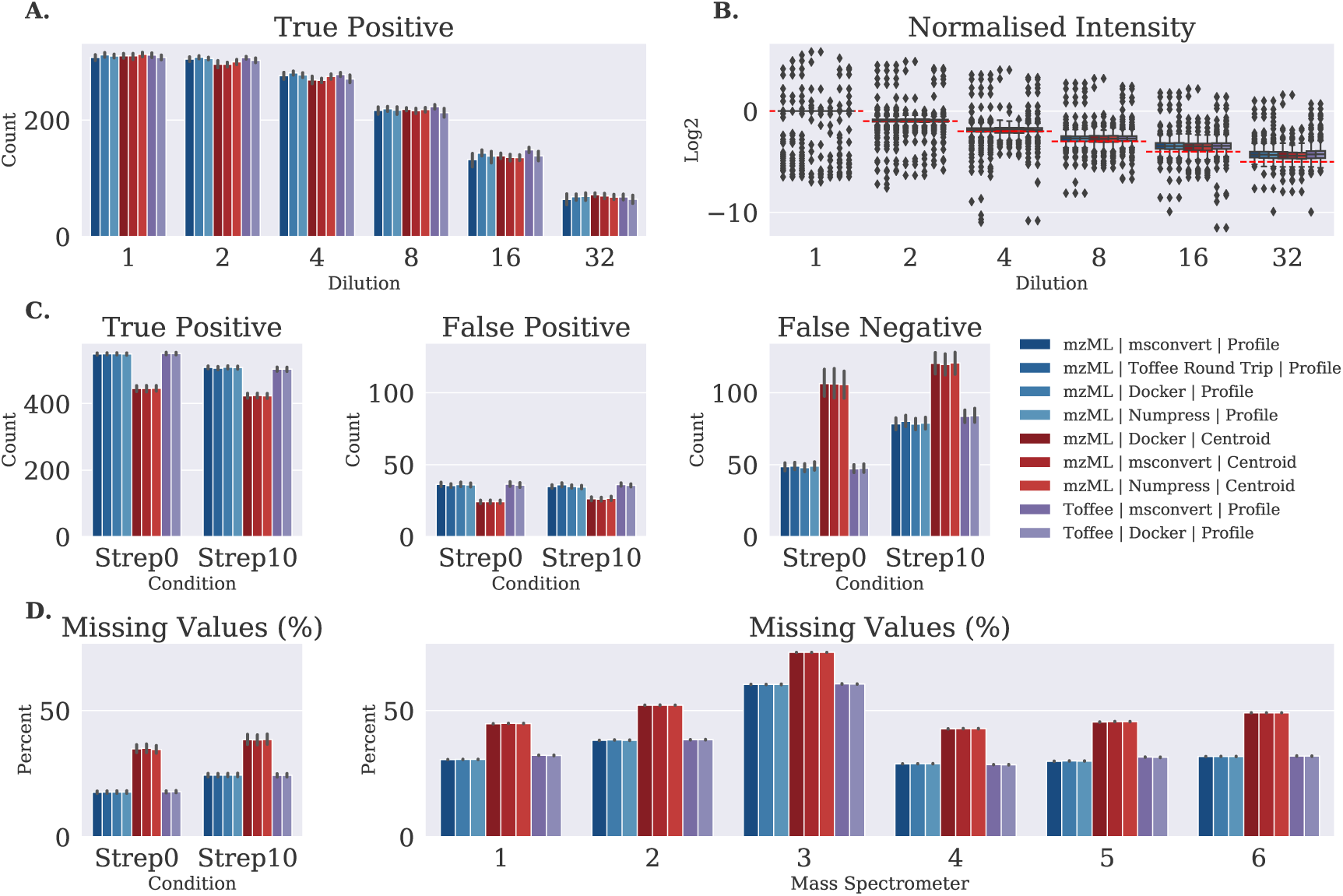
Analysis of four profile mzML file conversions, three centroid mzML file conversions, and two toffee file conversions. **A.** shows the number of true positive discoveries in the Swath Gold Standard data set and **B.** is their normalized intensity – from this data set there is essentially no difference between file conversion method. **C.** shows the confusion matrix for the TRIC manual validation data set clearly highlighting the degradation of centroiding during mzML creation. **D.** shows the percentage of missing values in the final peptide matrix, mimicking the false negative plot, for both the TRIC and ProCan90 data sets.

These results show there is no meaningful difference to the final quantified peptide results from OpenSWATH regardless of using profile data from ‘msconvert’ (with or without ‘msnumpress’), ‘Sciex Docker’, or toffee; however, there is a drop in performance when using centroided data.

### Deployment at scale

ProCan has moved to a toffee-based production pipeline since mid-2019. The program runs its analytics pipelines on a hybrid-cloud computational infrastructure with kubernetes orchestrating between on-premise and Amazon Web Services. By Production run times (not controlled for spectral library complexity or computational hardware) eliminating the need for mzML files, costs have decreased by an order of magnitude from a predicted $US 60,000 per month to $US 6,000 per month in year 5 of operation. In production, ProCan converts directly from the vendor file to toffee in a fully Dockerised workflow, and though file conversion is a once off, it is interesting to note that time taken to convert scales linearly with the size of the vendor file (see Figure 3a).

**Figure 3.**
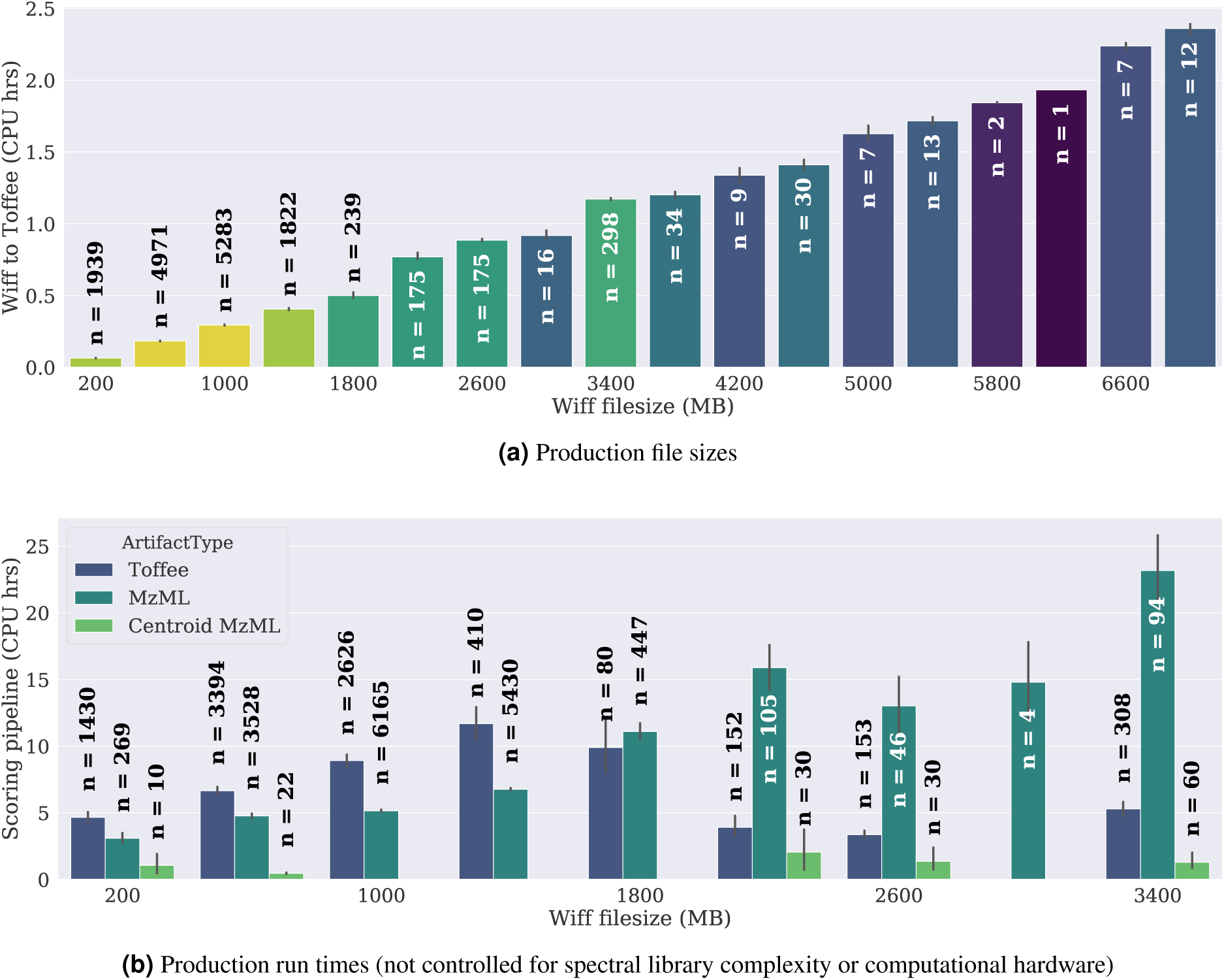
Timing of production *wiff-to-toffee* file conversions and OpenMSToffee / OpenSWATH runs for more than 10,000 mass spectrometer injections run at ProCan since mid-2017. As expected, CPU time scales roughly linearly with the size of the wiff file, however, exact timings will be dependent on the computational hardware available, in particular the I/O bandwidth.

Although Figure 3b does not control for the complexity of spectral library or the computational hardware (e.g. Amazon instance type), it is possible to observe processing time for toffee compared with mzML. Due to the architecture of OpenSWATH (in particular, challenges around thread safety), OpenMSToffee is by no means an optimum deployment of the technology and there is significant room for CPU performance improvements once const-correctness is addressed in OpenSWATH. For that reason, no attempt has been made to push toffee or OpenMSToffee into the upstream ‘openms’ repository.

### Novel uses

Having confirmed the OpenMSToffee pipeline is equivalent to the profile mzML/OpenSWATH pipeline, novel uses of toffee files can be explored.

#### In-silico dilution series

It is of critical importance to developing robust scientific software that one can isolate and test algorithms with controlled inputs and known expected outputs. By providing an efficient peptide-centric interface into the data, toffee allows algorithm developers to write tests of their implementation down to individual units of work while still retaining the complexity of real world data. This approach is fundamental to writing robust software^20^, for example, in high-energy physics, the US National Laboratories have developed the Tri-Lab test suite that pits algorithms against *toy* problems with analytic solutions^21^.

In the context of mass spectrometry, curating known input data is much more difficult due to the stochastic nature of the instruments, and of the subject under study. Often, validation experiments are based around injecting samples that contain a controlled dilution of a known group of peptides and technical replicates are performed to normalize the stochastic impact of the instrument. An alternative approach is to create data *in-silico* through analytic models^22, 23^; however, it is highly improbable that the noise artificially added to these models is an accurate reflection of data in reality.

Toffee offers us a new approach. By treating the data as triplets on a Cartesian grid, it becomes trivial to extract data from one toffee file, the foreground data, and add it to another toffee file, the background data, to create an entirely new toffee file. Referred to as an *in-silico dilution series*, data for specific peptides are extracted from the foreground file, scaled based on a *theoretical* dilution, and placed into the background file at a known retention time (see Figure 4).

**Figure 4.**
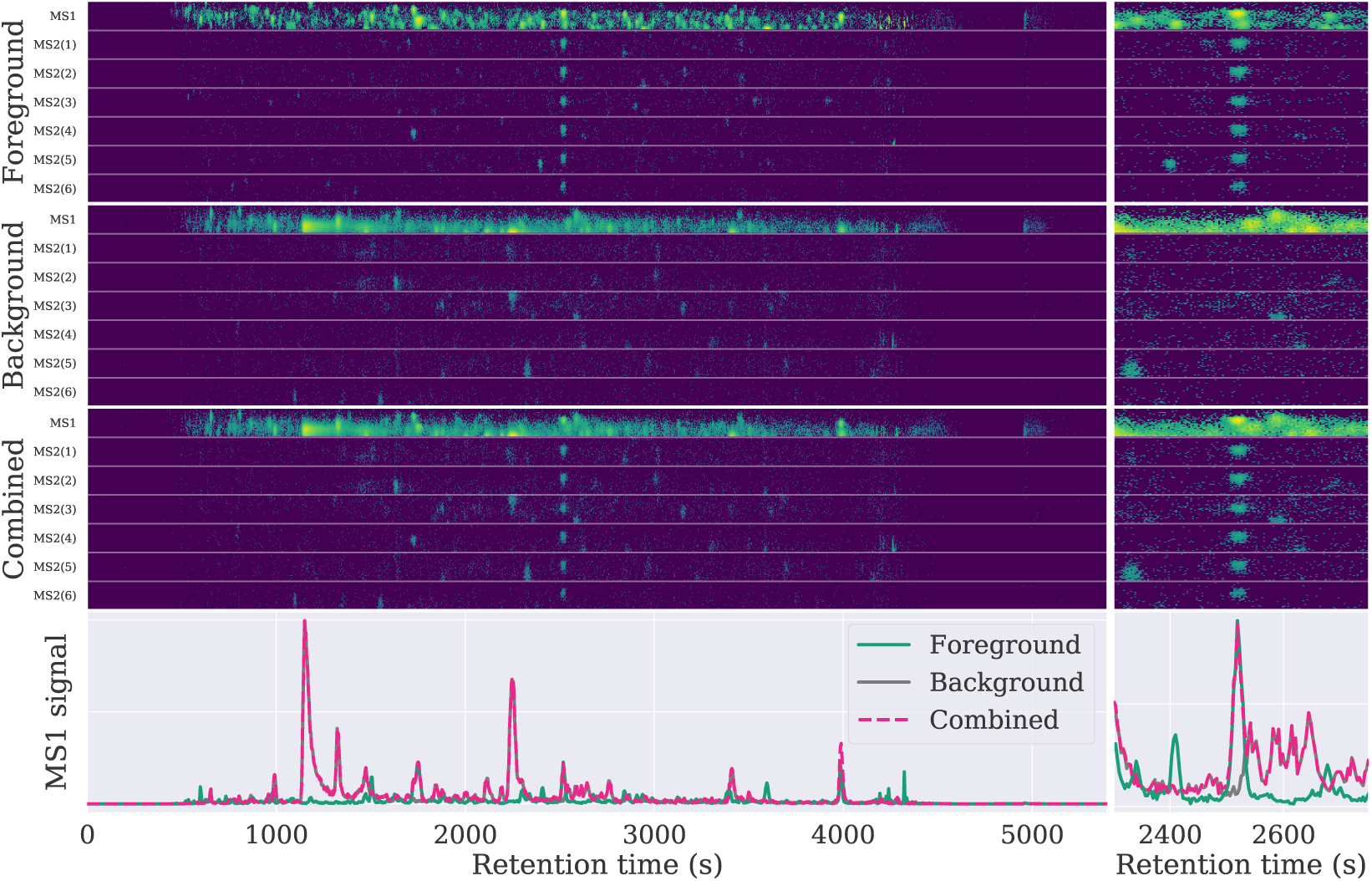
Combining data from a background file of *E.coli* and a foreground file of HEK293 for the peptide SKPGAAMVEMADGYAVDR charge 3. Shown in the top row are the HEK293 foreground data extracted for MS1 and the top six MS2 fragments, the middle row is the *E.coli* background data, and in the bottom row, the foreground HEK293 data (isolated at the peak retention time) has been added to the background with a theoretical dilution of 1.

In this study, two in-silico dilutions are constructed – one with a water background and another with an *E.coli* background – such that the impact of background noise can be assessed. Details of how this is done can be seen in Supplementary Material *in-silico-dilution.ipynb*. The files are then analyzed with two spectral libraries: a ‘Simple’ SRL containing just those peptides that were added in-silico, and a ‘Complex’ SRL that includes the in-silico peptides, plus an SRL derived from this *E.coli* sample. Comparing the results from these two SRLs neatly shows the impact of the *π*_0_ parameter discussed at length in Rosenberger et al.^17^

Figure 5A shows the normalized intensity quantified by the OpenMSToffee pipeline; as expected, the dilution curve of the water background is bound on the upper limit by the theoretical dilution (red horizontal lines) and the data that is below the theoretical limit is due to signal truncation at the lower limit of detection of the mass spectrometer (as set by the background file). In contrast, the *E.coli* dilution series often exceeds the theoretical limit and shows the OpenSWATH peak integration algorithms are at risk of quantifying noise by treating the extracted ion chromatogram as a one-dimensional signal rather than its two-dimensional reality. Figures 5B-5D show the confusion matrix where PQMs detected more than 10 seconds from the theoretical retention time are labelled as false positives, and PQMs correctly identified in the Simple SRL / Water Background for a given dilution and not detected in the file of interest are labelled as false negatives. From Figures 5B and 5C the complexity of the SRL seems unimportant at low dilutions and false positives remain around the expected FDR. At dilutions 4 and above, the more complex *E.coli* background makes it harder for PyProphet to distinguish between target and decoy peptides, leading to a more stringent FDR, and thus, a marked increase in the number of false negatives when compared to the water background (see Figure 5D).

**Figure 5.**
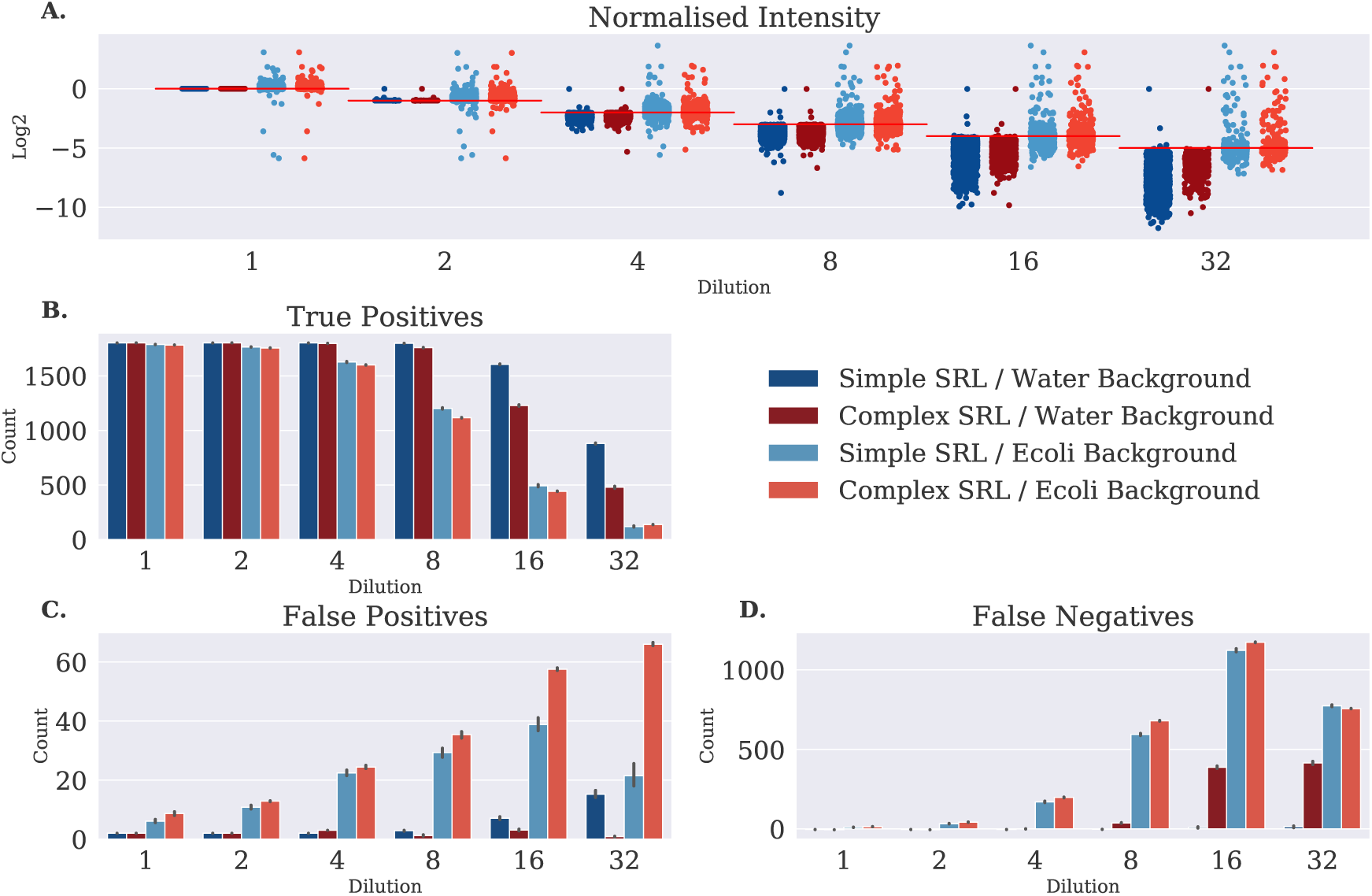
Analysis of the *in-silico* dilution series using simple and complex spectral libraries. **A.** shows the normalized intensity of each peptide query match against the expected value found in the most concentrated water background. **B.**–**D.** shows the confusion matrix found by comparing the retention time of the discovered peptide query match with the theoretical retention time at which the raw data was added. This clearly shows that the *E.coli* background leads to less true discoveries, and more false discoveries. Furthermore, FDR control with the complex spectral library leads to increased false negatives at high dilutions.

#### Re-Quantification of peaks

Through the many visualisations of toffee data, and appreciation for the TOF detector, one recognises that the data is roughly Gaussian in both retention time and m/z index space. By fitting an analytic model to the data, and using this model for *re-quantification* of peptide intensities, more accurate data can be obtained. Furthermore, by fixing the retention time of the analytic function for all fragments, it is also possible to deconvolute co-eluting peaks and count only the contribution from the peak of interest.

In a TOF mass analyzer, one can assume that individual ions obtain a distribution of kinetic energy that leads to an approximately normal distribution of data in m/z index space. Further, the elution profile of ions from the LC column can be approximated as a log-normal distribution that is skewed towards the left. These observations imply that numerical optimization can find the peak location, spread, and amplitude of the Gaussian functions for each fragment, *j*, through the following function:

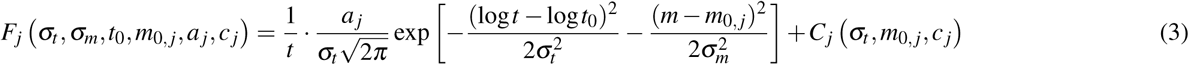

with chemical noise, for the MS1 signal only, defined as

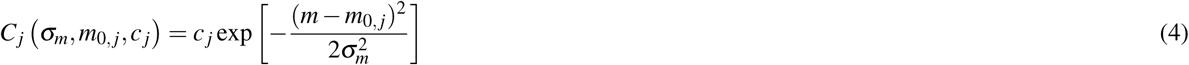

and the minimisation function

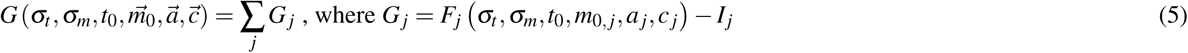

where *t* and *m* represent the retention time and 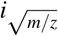 space respectively; *t*_0_ and 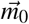 are the peak locations assuming the retention time for all fragments must be constant and the mass offset is allowed to be different for each one to account for calibration offsets; *σ*_*t*_, *σ*_*m*_ are the spread of each Gaussian; 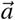 is the amplitude for each fragment; 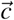 is the amplitude of chemical noise for each MS1 fragment; and *I*_*j*_ is the raw intensity data for a given fragment.

Figure 6 shows the result of applying this analytic model to more than 1,200 technical replicates selected from routine operation within ProCan. Interestingly, there is little change between the replicate correlation on the raw re-quantified data. However, it is possible to use the fit model parameters to perform an additional round of outlier removal. Specifically, the spread parameters *σ*_*t*_ and *σ*_*m*_, and the peak location parameters *t*_0_ and 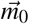 should be largely systematic features of the data. Peaks that were fit with parameters that were significantly different from their cohort are marked as false positives and dropped from the subsequent correlation calculation. In doing so, the correlation between technical replicates improves.

**Figure 6.**
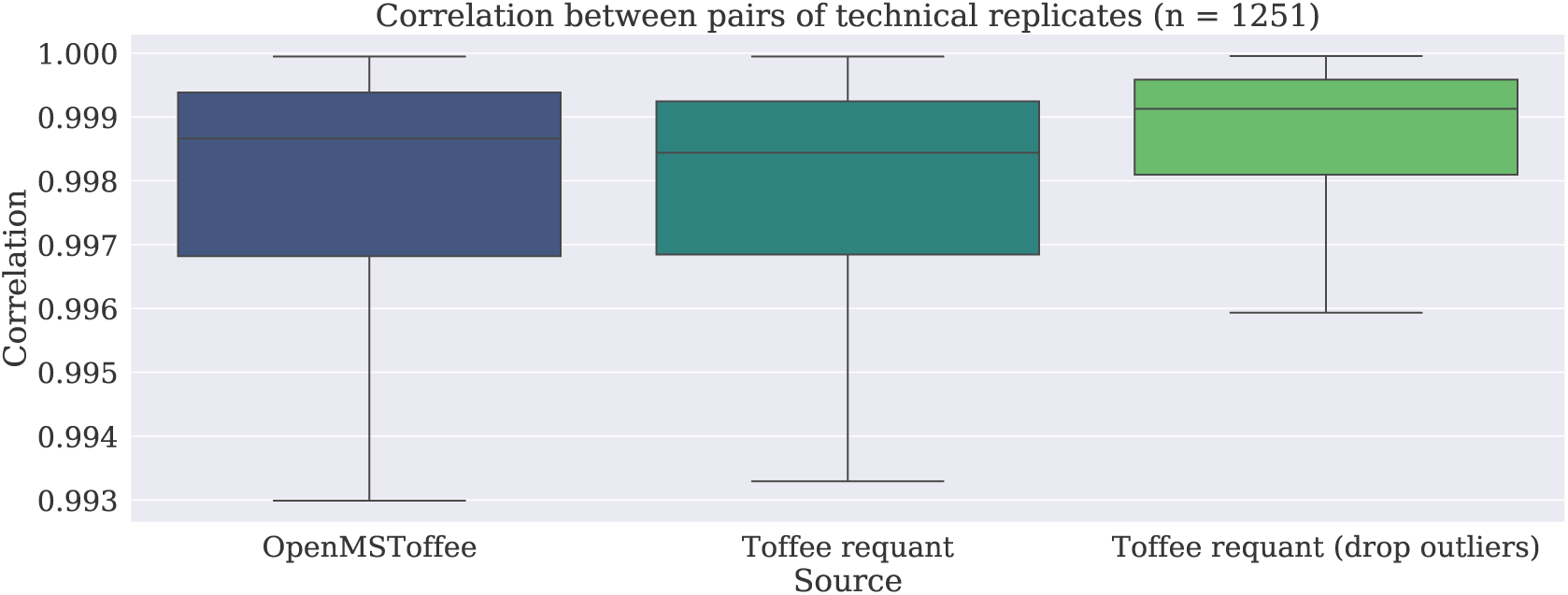
Correlation between peptide intensities of technical replicates before and after *re-quantification*. In this plot, correlation is calculated between the intensity of 1,251 technical replicates collected during routine production in ProCan. Using parameters found by fitting the analytic model to the data, potential outliers, or false discoveries, are detected and subsequently dropped from the analysis improve the correlation between replicates. Prior to outlier filtering, there is little difference between the re-quantified data and the original intensities recorded by OpenMSToffee both validating the analytic model chosen whilst indicating that there is not a significant burden in the data from co-eluting peptides.

#### Deep-learning pipeline proof-of-concept

While machine learning, and in particular deep learning, are permeating many facets of science, computer vision is the area where the technology is most mature. By treating toffee data as triplets on a Cartesian grid, and accepting a degree of mass approximation (<5 parts per million, see Supplementary Material *calculate_ims.ipynb*), it becomes trivial to extract data as a two-dimensional slice analogous to an image, and thus amenable to deep-learning based peptide identification strategies.

Figures 7 and 8 show raw two-dimensional data for two peptide query parameters extracted directly from a HEK293 file. Here, the red, green, and blue channels are filled with data extracted around the m/z of the precursor and product ions with offsets of 0, 1, and 2 times the isotopic carbon-12 / carbon-13 mass difference. Using OpenSWATH results from ProCan90, it is possible to develop a significant labelled training dataset suitable for training an object detection convolutional neural network. For this proof of concept, a single shot detector (SSD) architecture was trained with a ResNet50 backbone on the Amazon SageMaker platform (see Supplementary Material *raptor-part0-transfer_from_resnet50.ipynb*). Figure 8 shows inference results for a random sample of target peptides from the holdout set where a-d are successfully detected peptides and e-f are false negatives.

**Figure 7.**
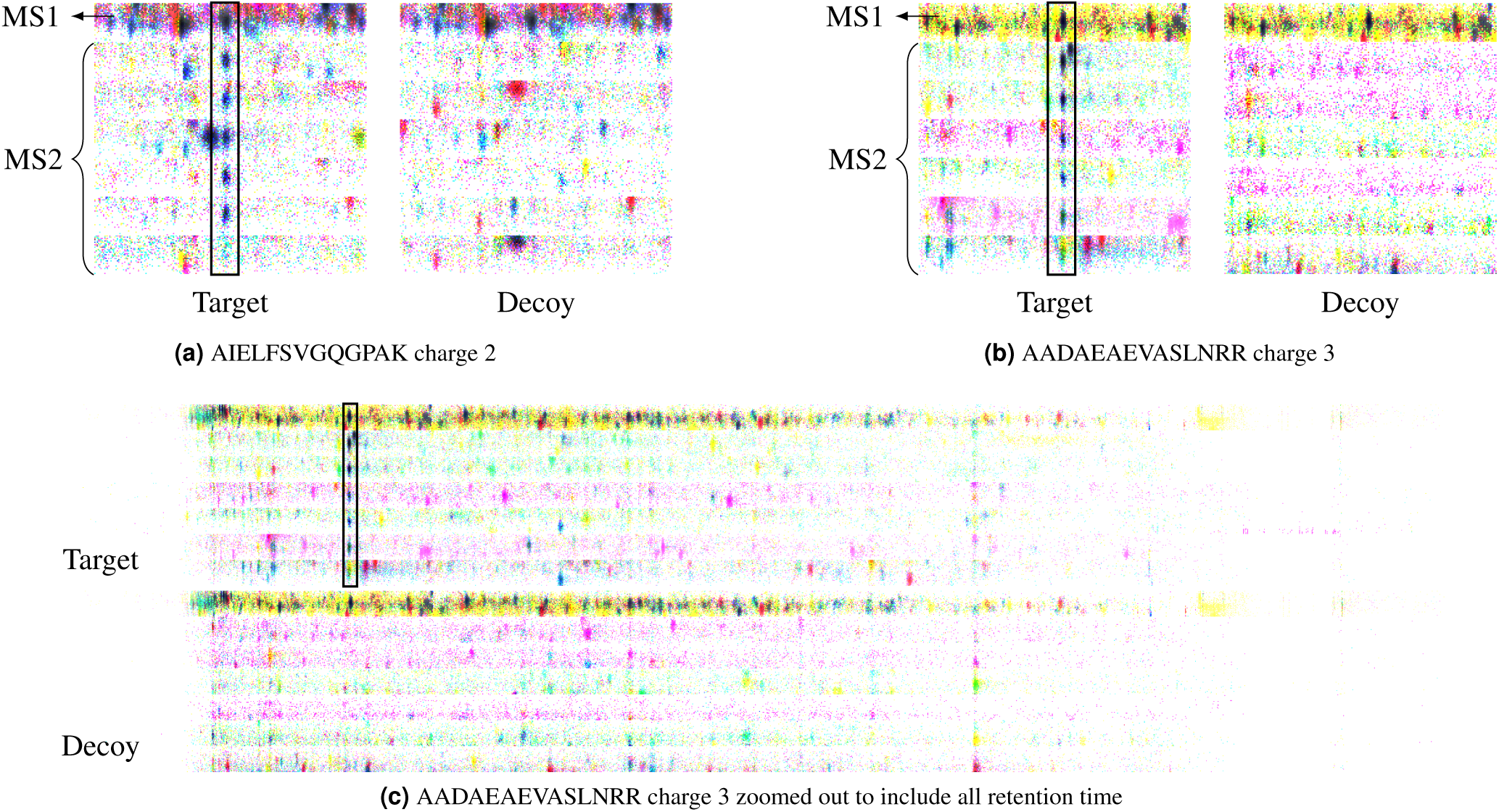
By accepting a mass accuracy loss of less than 5 parts per million, toffee data can be efficiently accessed in a way that is amenable for modern computer vision algorithms. For example, the images here are produced by stacking data from the mono-isotope, and isotopes 1 and 2, into the red, green and blue channels of an image respectively. The left panel shows target and corresponding data for AIELFSVGQGPAK charge 2, and the right panel is for AADAEAEVASLNRR charge 3. The top two-dimensional horizontal slice of each of these images is from the precursor ion (or MS1) data, while the remaining slices are from the six product ion (or MS2) data as specified in the spectral library. Images **a** and **b** are zoomed to the retention time location of the peptide, however as shown in **c**, it is possible to keep the full retention time range if one desires.

**Figure 8.**
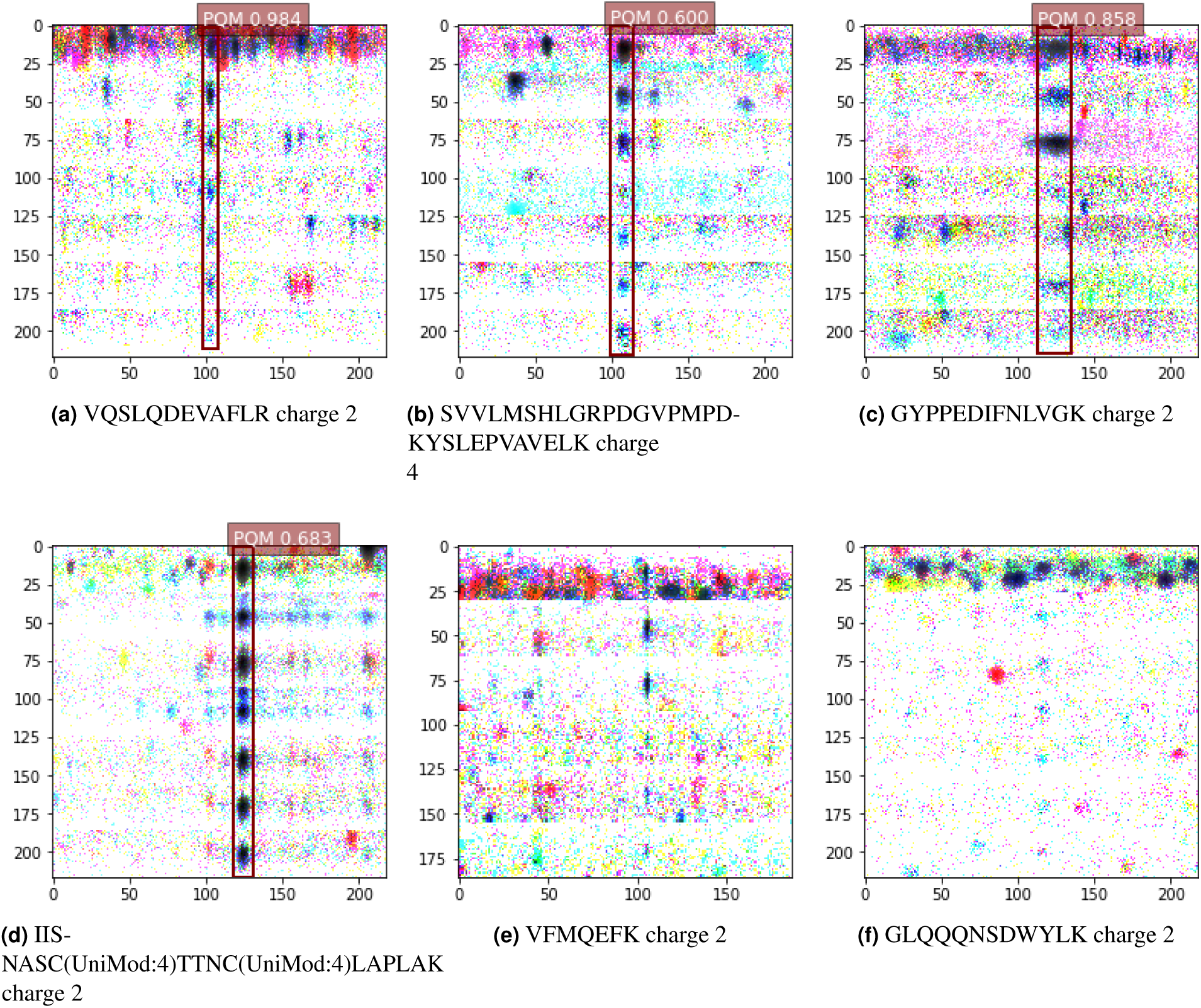
Inference of peptide location using a single-shot detector deep-learning algorithm. These very early prototype results confirm that modern computer vision algorithms are highly amenable to use for peptide-centric analysis of DIA-MS data. False-negative results can be seen in **e** and **f**, where the algorithm fails to detect and score a bounding box around the expected retention time. *(Note: UniMod:4 denotes Carbamidomethyl modification)*

Clearly, this proof of concept is not yet at the performance of state-of-the-art SWATH-DIA analysis tools. However, the results are encouraging and give hope that an analyis pipeline that does not require an experimental spectral library may be possible in the not too distant future.

## Discussion

In summary, the challenges of file size and data access with current open formats for data independent acquisition mass spectrometry are acute when a scientific program needs to operate at biobank-scale. These challenges significantly increase the cost and complexity of data management and analysis, and hold back the progress of writing efficient algorithms that are routinely tested on real-world data. Toffee aims to address these issues by taking a first-principles approach to understanding the raw data, and translating those findings into a best-practice software library. Code is released with an MIT license, python packages should be easily installed for MacOS and Linux using ‘conda’, both toffee and an exemplar implementation wrapping OpenSWATH are available via version-controlled Docker images, and all analyses performed in this paper are available as Jupyter notebooks.

## Methods

### Open Science

To the extent possible, this work aims to have full and automated reproducibility. The code is released with an MIT license and makes use of the following community tools and technologies: pyteomics^24, 25^; psims^26^; msconvert^13, 14^; OpenSWATH^10^; PyProphet^17^; numpy^27^; scipy^28^; pandas^29^; matplotlib^30^; plotly^31^; HDF5^7^; h5py^32^; Eigen^33^; Project Jupyter^34^; and Docker^35^.

#### Data sets

Three publicly available data sets are used in the current work: Swath Gold Standard and TRIC data available from PeptideAtlas raw data repository with accession number PASS00289^10^ and PASS00788^11^, respectively, and the ProCan90 dataset can be obtained from the PRIDE archive under the identifier PXD011093^12^.

#### Software and Analysis

- Code repositories
  – https://bitbucket.org/cmriprocan/toffee
  – https://bitbucket.org/cmriprocan/openms-toffee
  – https://bitbucket.org/cmriprocan/dia-test-data – a resource of small DIA-MS files that are used in continuous integration regression tests
  – https://bitbucket.org/cmriprocan/openms-toffee-paper – all analyses produced for this paper
  – https://bitbucket.org/cmriprocan/openms – a versioned Docker file for creating the base OpenMS Docker image
- Documentation
  – https://toffee.readthedocs.io
  – https://openms-toffee.readthedocs.io
- Docker images
  – `cmriprocan/toffee`
  – `cmriprocan/openms-toffee`
  – `cmriprocan/openms`
- Python conda library: `conda install -c cmriprocan -c plotly toffee`

### File conversion

Original Sciex wiff files were converted to various forms of mzML, mz5, and toffee files. While conversion methods are provided here for reference, a self-contained script is included in the Supplementary Material *convert_mzml.py*.

**mzML | msconvert | Profile:** Conversion of wiff to mzML using msconvert version 3.0.18304 in a windows environment

~~~
msconvert . exe ––mzML –z \
  ––outfile $ {mzml_fname} \
  $ {wiff_ fname}
~~~

**mzML | msconvert | Centroid:** Conversion of wiff to mzML using msconvert version 3.0.18304 in a windows environment, and applying vendor peak picking to centroid the data

~~~
msconvert . exe ––mzML –z \
  ––filter “peakPicking vendor” \
  ––outfile $ {mzml_fname} \
  $ {wiff_ fname}
~~~

**mzML | Numpress | Profile:** Conversion of wiff to mzML using msconvert version 3.0.18304 in a windows environment, and applying the Numpress compression

~~~
msconvert . exe ––mzML –z \
  ––numpres s Linear \
  ––outfile $ {mzml_fname} \
  $ {wiff_ fname}
~~~

**mzML | Numpress | Centroid:** Conversion of wiff to mzML using msconvert version 3.0.18304 in a windows environment, and applying the Numpress compression algorithm and vendor peak picking to centroid the data

~~~
msconvert . exe ––mzML–z \
  ––numpressLinear \
  ––filter “peakPicking vendor” \
  ––outfile $ {mzml_fname} \
  $ {wiff_ fname}
~~~

**mz5 | msconvert | Profile:** Conversion of wiff to mz5 using the Dockerised version of msconvert `chambm/pwiz-skyline-i-agree-`

~~~
wine msconvert z mz5 \
  ––outfile $ {mz5_fname} \
  $ {wiff_ fname}
~~~

**mzML | Docker | Profile:** Conversion of wiff to mzML using the publicly available Sciex Docker image `sciex/wiffconverter:0.9`

~~~
 mono \
  / usr / local / bin / sciex / wiffconverter / OneOmics . WiffConverter . exe \
  WIFF $ {wiff_ fname} \
  – profile MZML $ {mzml_fname} \
  –– zlib –– index
~~~

**mzML | Docker | Centroid:** Conversion of wiff to mzML using the publicly available Sciex Docker image `sciex/wiffconverter:0` and applying the included peak picking algorithm

~~~
mono \
  / usr / local / bin / sciex / wiffconverter / OneOmics . WiffConverter . exe \
  WIFF $ {wiff_ fname} \
  – centroid MZML $ {mzml_fname} \
  –– zlib –– index
~~~

**Toffee | msconvert | Profile:** Conversion of the msconvert profile mzML to toffee using the toffee Docker image `cmriprocan/toffee:0.12.16` ^37^

~~~
mzml _ to _ toffee $ {mzml_fname} $ {tof _ fname}
~~~

**Toffee | Docker | Profile:** Conversion of the Sciex Docker profile mzML to toffee using the toffee Docker image `cmriprocan/toffee:0.12.16` ^37^

~~~
mzml _ to _ toffee $ {mzml_fname} $ {tof _ fname}
~~~

**mzML | Toffee Round Trip | Profile:** Using the toffee file created using the Sciex Docker profile mzML, convert back to mzML using the toffee Docker image `cmriprocan/toffee:0.12.16`^37^

~~~
toffee _ to _ mzml $ {tof _ fname} $ {mzml_fname}
~~~

### DIA pipeline

All analysis for raw data was completed using the Docker image `cmriprocan/openms-toffee:0.13.12.dev`^38^ and a complete python script for marshalling analysis is included in the Supplementary Material *pipeline.py*. In short, the current best-practice OpenSWATH workflow is used whereby the spectral library is provided in SQLite format, scores are saved to an `osw` result file, and PyProphet (version 2.0.4; git hash d35a53af86131e7c4eb57bbb09be8935a1f30c70) FDR control is applied at the peak-group, peptide, and protein levels. For the latter two, scores are calculated across the full cohort of the experiment (i.e. the ‘global’ context is used); in a departure from the defaults, the `-parametric` model is used as it was found to be less conservative on the small SRLs used in this study. mzML files are analyzed using a modified version of OpenMS v2.4.0 (`CMRI-ProCan/OpenMS` version `CMRI-ProCan-v1.1.2`^39^). This same version is used as the basis for the code in the OpenMSToffee bitbucket repository^18^, a wrapper around OpenMS that enables toffee files to be used in conjunction with the algorithms of OpenSWATH. Furthermore, OpenMSToffee has a series of tests that ensure regressions are not introduced to OpenSWATH scoring algorithms when the version of OpenMS is updated. Finally, `OpenMSToffeeWorkflow` is used as a drop-in replacement for `OpenSwathWorkflow` when analyzing toffee files.

### Toffee file format

#### Time of Flight (TOF) m/z transfer function

The mass analyzer in a TOF mass spectrometer is a digital sensor that samples at a constant frequency. It effectively measures the time taken for a charged ion to move a known distance (*d*) through an electric field of known strength (*U*). Thus, measurements are made at integer multiples (*i*) of a constant time interval (Δ*t*), such that

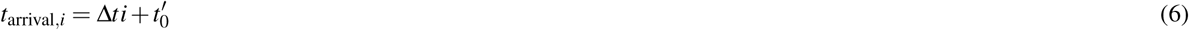

where 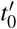 is an offset applied such that *t*_arrival,0_ makes sense. This time of arrival is related to the mass over charge of the ion through the electric potential energy, *E*_*p*_ = *zU*, and kinetic energy, 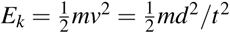, such that

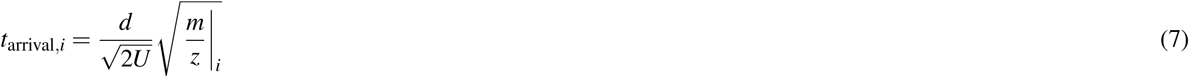

With this in mind, it becomes possible to convert mass over charge values to an integer index through the following,

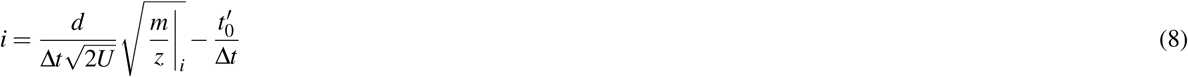

or equivalently,

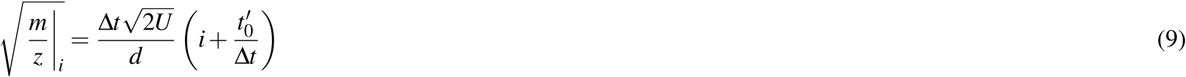

Then introducing the intrinsic mass spacing parameters, *α, β*, and *γ*

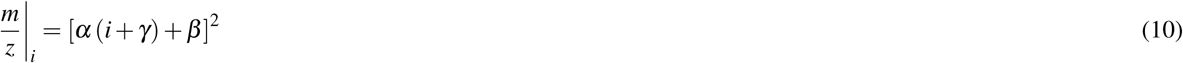

where

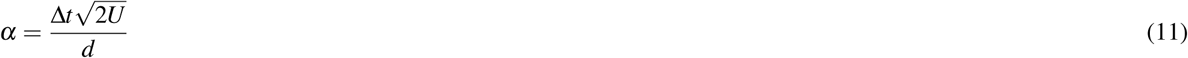

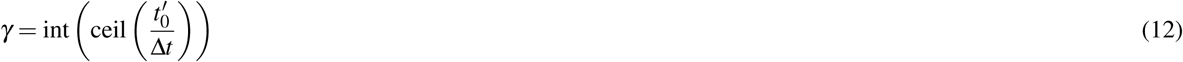

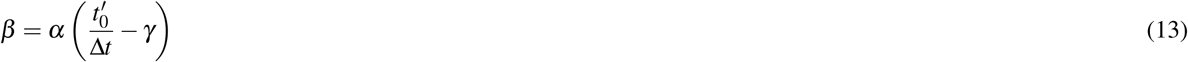

#### HDF5 structure

The toffee HDF5 file is structured as follows:

**attr:CREATED_BY_LIBRARY_VERSION (str)** the version of the toffee library used to create the file

**attr:FILE_FORMAT_MAJOR_VERSION (int)** the major version of the toffee file format of this file

**attr:FILE_FORMAT_MINOR_VERSION (int)** the major version of the toffee file format of this file

**attr:IMSType (str)** the IMS type of the file. Currently, toffee is tested on “TOF”, however, infrastructure is in place for “Orbitrap”

**attr:metadataXML (int)** An mzML style XML header that mimics the header of a normal mzML file

**group:ms1** A group representing the MS1 data

**attr:IMSAlpha (double)** the median IMS *α* parameter for this window
**attr:IMSBeta (double)** the median IMS *β* parameter for this window
**attr:IMSGamma (int)** the IMS *γ* parameter for this window
**attr:firstScanRetentionTimeOffset (double)** the first retention time for scans in this window
**attr:scanCycleTime (double)** the cycle time between scans for this window – this is expected (but not enforced) to be constant for all windows in the toffee file
**attr:precursorLower (double)** the lower precursor m/z bound of this window (−1 / ignored for MS1)
**attr:precursorCenter (double)** the center of precursor m/z bounds of this window (−1 / ignored for MS1)
**attr:precursorUpper (double)** the upper precursor m/z bound of this window (−1 / ignored for MS1)
**dataset:IMSAlphaPerScan (double)** the IMS *α* parameter for each scan of this window
**dataset:IMSBetaPerScan (double)** the IMS *β* parameter for each scan of this window
**dataset:retentionTimeIdx (unsigned int32)** the equivalent of the ‘index pointer’ of a compressed sparse row matrix.
For toffee, there is one entry for each scan in the window and the value represents the **last** index of the `imsCoord` and `intensity` that is valid for this scan. For example `start = retentionTimeIdx[i - 1]; end = retentionTimeIdx[i]; scan_i_intensity = intensity[start:end]`.
**dataset:imsCoord (unsigned int32)** the IMS m/z coordinate vector
**dataset:intensity (unsigned int32)** the intensity vector

**group:ms2-(001, …, n)** Groups with the same format as ‘ms1’ that represent the MS2 data

## Supporting information

Jupyter notebooks

## Acknowledgements

I would like to acknowledge Keith Ashman, Peter Hains, Phil Robinson, and Akila Seneviratne for their help in understanding the structure of TOF DIA-MS data; David Clarke, Sean Peters, and Akila Seneviratne for feedback and suggestions for the analyses conducted in this work; and Michael Daussman, Michael Hecker, Sean Peters, and Max Witmann for the tireless work of automated deployment of DIA-MS pipelines at scales beyond 10,000 mass spectrometer injections. ProCan is supported by the Australian Cancer Research Foundation, Cancer Institute New South Wales (NSW) (2017/TPG001, REG171150), NSW Ministry of Health (CMP-01), University of Sydney, National Breast Cancer Foundation (IIRS-18-164), Cancer Council NSW (IG 18-01), Ian Potter Foundation, the Medical Research Futures Fund (MRFF-PD), and the National Health and Medical Research Council of Australia (GNT1047070; GNT1170739). This project was supported by equipment grants from the Australian Cancer Research Foundation (ACRF), Australia, the Cancer Institute NSW (CINSW) as a Research Equipment Grant (TPG), The Ian Potter Foundation, and Perpetual IMPACT Philanthropy.

## Author contributions statement

BT conceived the idea, executed the work, and wrote the manuscript.

## Additional Information

The authors declare no competing interests.

## References

1. Martens, L. et al. mzml—a community standard for mass spectrometry data. Mol. & Cell. Proteomics 10, DOI: 10.1074/mcp.R110.000133 (2011). https://www.mcponline.org/content/10/1/R110.000133.full.pdf.

2. Wilhelm, M., Kirchner, M., Steen, J. A. J. & Steen, H. mz5: Space- and time-efficient storage of mass spectrometry data sets. Mol. & Cell. Proteomics 11, DOI: 10.1074/mcp.O111.011379 (2012). https://www.mcponline.org/content/11/1/O111.011379.full.pdf.

3. Bouyssié, D. et al. mzDB: A File Format Using Multiple Indexing Strategies for the Efficient Analysis of Large LC-MS/MS and SWATH-MS Data Sets. Mol. & Cell. Proteomics 14, 771–781, DOI: 10.1074/mcp.O114.039115 (2015).

4. Nasso, S. et al. An optimized data structure for high-throughput 3D proteomics data: mzRTree. J. Proteomics 73, 1176–1182, DOI: 10.1016/j.jprot.2010.02.006 (2010). 1002.3724v2.

5. Handy, K., Rosen, J., Gillan, A. & Smith, R. Fast, axis-agnostic, dynamically summarized storage and retrieval for mass spectrometry data. PLoS ONE 12, 1–14, DOI: 10.1371/journal.pone.0188059 (2017).

6. Guttman, A. R-trees: A dynamic index structure for spatial searching. In INTERNATIONAL CONFERENCE ON MANAGEMENT OF DATA, 47–57 (ACM, 1984).

7. The HDF Group. Hierarchical Data Format, version 5 (1997–2019). [Online; accessed 4-June-2019].

8. Schneider, L. Mass spectral data processing. Tech. Rep., Veritomyx (2016). DOI: 10.13140/RG.2.2.26279.75684.

9. Wikipedia contributors. Sparse matrix — Wikipedia, the free encyclopedia. https://en.wikipedia.org/w/index.php?title=Sparse_matrix&oldid=892846660 (2019). [Online; accessed 18-April-2019].

10. Röst, H. L. et al. OpenSWATH enables automated, targeted analysis of data-independent acquisition MS data. Nat. Biotechnol. 32, 219–223, DOI: 10.1038/nbt.2841 (2014).

11. Röst, H. L. et al. Tric: an automated alignment strategy for reproducible protein quantification in targeted proteomics. Nat. methods 13, 777—783, DOI: 10.1038/nmeth.3954 (2016).

12. Peters, S., Hains, P. G., Lucas, N., Robinson, P. J. & Tully, B. A case study and methodology for openswath parameter optimization using the procan90 data set and 45,810 computational analysis runs. J. Proteome Res. 18, 1019–1031, DOI: 10.1021/acs.jproteome.8b00709 (2019). PMID: 30652484, https://doi.org/10.1021/acs.jproteome.8b00709.

13. Kessner, D., Agus, D., Chambers, M., Mallick, P. & Burke, R. ProteoWizard: open source software for rapid proteomics tools development. Bioinformatics 24, 2534–2536, DOI: 10.1093/bioinformatics/btn323 (2008). http://oup.prod.sis.lan/bioinformatics/article-pdf/24/21/2534/16882584/btn323.pdf.

14. Chambers, M. C. et al. A cross-platform toolkit for mass spectrometry and proteomics. Nat. Biotechnol. 30, 918–920, DOI: 10.1038/nbt.2377 (2012).

15. Teleman, J. et al. Numerical compression schemes for proteomics mass spectrometry data. Mol. & Cell. Proteomics 13, 1537–1542, DOI: 10.1074/mcp.O114.037879 (2014). https://www.mcponline.org/content/13/6/1537.full.pdf.

16. Sciex. Docker image: sciex/wiffconverter:0.9. https://hub.docker.com/r/sciex/wiffconverter (2018). [Online; accessed 18-April-2019].

17. Rosenberger, G. et al. Statistical control of peptide and protein error rates in large-scale targeted data-independent acquisition analyses. Nat. Methods 14, 921 (2017).

18. CMRI ProCan Software Engineering. Bitbucket code repository for openms-toffee. https://bitbucket.org/cmriprocan/openms-toffee (2019). [Online; accessed 18-April-2019].

19. Brett Tully. Analysis code for openms-toffee paper. https://bitbucket.org/cmriprocan/openms-toffee-paper (2019). [Online; accessed 18-April-2019].

20. Wilson, G. et al. Best practices for scientific computing. PLOS Biol. 12, 1–7, DOI: 10.1371/journal.pbio.1001745 (2014).

21. J. S. Brock, W. J. R. S. B. C. W. P. K. T. G. T., J. R. Kamm. Verification test suite for physics simulation codes. Tech. Rep., Lawrence Livermore National Laboratory (2006). [Online; accessed 18-April-2019].

22. Bielow, C., Aiche, S., Andreotti, S. & Reinert, K. Mssimulator: Simulation of mass spectrometry data. J. Proteome Res. 10, 2922–2929, DOI: 10.1021/pr200155f (2011). PMID: 21526843, https://doi.org/10.1021/pr200155f.

23. Awan, M. G. & Saeed, F. Mass-simulator: A highly configurable simulator for generating ms/ms datasets for benchmarking of proteomics algorithms. PROTEOMICS 18, 1800206, DOI: 10.1002/pmic.201800206 (2018). https://onlinelibrary.wiley.com/doi/pdf/10.1002/pmic.201800206.

24. Goloborodko, A. A., Levitsky, L. I., Ivanov, M. V. & Gorshkov, M. V. Pyteomics—a Python Framework for Exploratory Data Analysis and Rapid Software Prototyping in Proteomics. J. The Am. Soc. for Mass Spectrom. 24, 301–304, DOI: 10.1007/s13361-012-0516-6 (2013).

25. Levitsky, L. I., Klein, J. A., Ivanov, M. V. & Gorshkov, M. V. Pyteomics 4.0: Five Years of Development of a Python Proteomics Framework. J. Proteome Res. 18, 709–714, DOI: 10.1021/acs.jproteome.8b00717 (2019).

26. Klein, J. & Zaia, J. psims - A Declarative Writer for mzML and mzIdentML for Python. Mol. & cellular proteomics : MCP 18, 571–575, DOI: 10.1074/mcp.RP118.001070 (2019).

27. van der Walt, S., Colbert, S. C. & Varoquaux, G. The NumPy Array: A Structure for Efficient Numerical Computation. Comput. Sci. & Eng. 13, 22–30, DOI: 10.1109/MCSE.2011.37 (2011).

28. Jones, E., Oliphant, T., Peterson, P. et al. SciPy: Open source scientific tools for Python (2001–2019). [Online; accessed 4-June-2019].

29. Numfocus. Python data analysis library – pandas: Python data analysis library. https://pandas.pydata.org/ (2018). [Online; accessed 5-September-2018].

30. Hunter, J. D. Matplotlib: A 2D Graphics Environment. Comput. Sci. & Eng. 9, 90–95, DOI: 10.1109/MCSE.2007.55 (2007).

31. Inc., P. T. Collaborative data science (2015). [Online; accessed 4-June-2019].

32. Collette, A. Python and HDF5 (O’Reilly, 2013).

33. Guennebaud, G., Jacob, B. et al. Eigen v3. http://eigen.tuxfamily.org (2010).

34. Project Jupyter. Project jupyter | home. http://jupyter.org/ (2018). [Online; accessed 5-September-2018].

35. Docker. Docker – build, ship, and run any app, anywhere. https://www.docker.com/ (2018). [Online; accessed 5-September-2018].

36. Matt Chambers. chambm/pwiz-skyline-i-agree-to-the-vendor-licenses:3.0.19073-85be84641. https://hub.docker.com/r/chambm/pwiz-skyline-i-agree-to-the-vendor-licenses (2019). [Online; accessed 18-April-2019].

37. CMRI ProCan Software Engineering. Docker image: cmriprocan/toffee:0.12.16. https://hub.docker.com/r/cmriprocan/toffee (2019). [Online; accessed 18-April-2019].

38. CMRI ProCan Software Engineering. Docker image: cmriprocan/openms-toffee:0.13.12.dev. https://hub.docker.com/r/cmriprocan/openms-toffee (2019). [Online; accessed 18-April-2019].

39. CMRI ProCan Software Engineering. Openms fork. https://github.com/CMRI-procan/OpenMS (2018). [Online; accessed 5-September-2018].

