## Supplementary material for "Toffee – a highly efficient, lossless file format for DIA-MS": Jupyter notebooks: analysis-error.ipynb

```
# %% [code]
%load_ext autoreload
%autoreload 2

# %% [code]
import h5py
import os

import matplotlib.pyplot as plt

import numpy as np

from tqdm import tqdm

import seaborn as sns

import pandas as pd

import xarray as xr

sns.set(
    palette='husl',
    font_scale=2,
)

# %% [code]
def plot_ppm_ax(w, ax):
    np.random.seed(0)
    converted = w['converted'][()]
    original = w['original'][()]
    rt = w['retentionTime'][()]
    ppm = 1e6 * (converted - original) / original
    mask = np.random.choice(original.shape[0], min(5000, original.shape[0]))
    max_rt = rt.max()
    color = np.array([plt.cm.viridis(r / max_rt) for r in rt[mask]])
    ax.scatter(x=original[mask], y=ppm[mask], s=4, c=color, alpha=0.5)
    ax.axhline(np.median(ppm), c='maroon', ls='--', alpha=0.5)

def plot_ppm(basename):
    fname_old_a = f'{basename}.old_a.err.h5'
    fname_a = f'{basename}.a.err.h5'
    fname_e = f'{basename}.e.err.h5'
    assert os.path.isfile(fname_old_a)
    
    # get windows to plot
    with h5py.File(fname_old_a, 'r') as old_a:
        windows = list(old_a.keys())
        windows = [
            windows[0],
            windows[1],
            windows[len(windows) // 2],
            windows[-1],
        ]
        
    fnames = [
        fname_a,
        fname_e,
        fname_old_a,
    ]
    ncols = len(fnames)
    nrows = len(windows)
    fig, axes = plt.subplots(nrows=nrows, ncols=ncols, figsize=(5 * ncols, 5 * nrows), sharey=True, sharex=True)
    
    for window_name, axe in tqdm(zip(windows, axes)):
        for fname, ax in zip(fnames, axe):
            if not os.path.isfile(fname):
                continue
            with h5py.File(fname, 'r') as f:
                plot_ppm_ax(f[window_name], ax)
    
    for window_name, ax in zip(windows, axes[:, 0]):
        ax.set_ylabel(f'{window_name.upper()} Error (ppm)')
    
    fig.tight_layout()

# %% [code]
plot_ppm(basename='results/roest2016/Strep0Repl1_R01_SW')

# %% [code]
plot_ppm(basename='results/procan90/ProCan90-M01-01')

# %% [code]
plot_ppm(basename='results/sgs/napedro_l120417_002_sw')

# %% [code]
```
