## Supplementary material for "Toffee – a highly efficient, lossless file format for DIA-MS": Jupyter notebooks: analysis-profile.ipynb

```
# %% [markdown]
# Comparing methods for creating profile mzML

We have typically used `msconvert` for creating the profile mzML files that are passed into the OpenSwath workflow. This requires a special windows machine to sit outside of our Kubernetes cluster and is a painful departure from a pure automated workflow. With the recent discovery of a Sciex Docker container for converting wiff files, we would like to compare that the data extracted is the same.

The files produced are different sizes, but with the following shows that the m/z and intensities are exactly the same, and the retention time differs only in that one stores minutes and the other stores seconds, so there is a truncation error in the conversion. In saying this, there is a difference in the spectra that is stored, in that `msconvert` "zero pads" the peaks -- i.e. zero intensities are added between non-consecutive m/z non-zero intensities.

# %% [code]
%load_ext autoreload
%autoreload 2

# %% [code]
import h5py

import matplotlib.pyplot as plt

import numpy as np

from tqdm import tqdm_notebook

import seaborn as sns

import pandas as pd

import xarray as xr

sns.set(
    palette='husl',
    font_scale=2,
)

# %% [code]
results = []
fname_a = 'raw_data/procan90/ProCan90-M03-01.a.ms2-050.spech5'
fname_b = 'raw_data/procan90/ProCan90-M03-01.b.ms2-050.spech5'
fname_e = 'raw_data/procan90/ProCan90-M03-01.e.ms2-050.spech5'
with h5py.File(fname_a, 'r') as raw_a, h5py.File(fname_b, 'r') as raw_b, h5py.File(fname_e, 'r') as raw_e:
    for window_name in raw_a.keys():
        w_a = raw_a[window_name]
        w_b = raw_b[window_name]
        w_e = raw_e[window_name]
        for scan_name in tqdm_notebook(w_a.keys()):
            s_a = w_a[scan_name]
            s_b = w_b[scan_name]
            s_e = w_e[scan_name]
            
            runs = [
                ('AB', s_a, s_b),
                ('AE', s_a, s_e),
                ('EB', s_e, s_b),
            ]
            
            for key, a, b in runs:
                delta_rt = a.attrs.get('RETENTION_TIME') - b.attrs.get('RETENTION_TIME')
                delta_i = a['INTENSITY'][()] - b['INTENSITY'][()]
                mz_a = a['MASS_OVER_CHARGE'][()]
                mz_b = b['MASS_OVER_CHARGE'][()]
                delta_mz = mz_a - mz_b
                ppm = 1e6 * delta_mz / mz_a
                r = {
                    'Window': f'{window_name}/{key}',
                    'Scan': scan_name,
                    'DeltaRetentionTime': np.sum(np.abs(delta_rt)),
                    'DeltaMassOverCharge': np.sum(np.abs(delta_mz)),
                    'Max_PPM': np.max(ppm),
                    'Min_PPM': np.min(ppm),
                    'Median_PPM': np.median(ppm),
                    'DeltaIntensity': np.sum(delta_i),
                }
                results.append(r)
results = pd.DataFrame(results)

# %% [code]
results.groupby('Window').describe().T

# %% [code]
```
