## Supplementary material for "Toffee – a highly efficient, lossless file format for DIA-MS": Jupyter notebooks: analysis.ipynb

```
# %% [code]
%load_ext autoreload
%autoreload 2

# %% [code]
import os
from collections import OrderedDict

import matplotlib as mpl
import matplotlib.pyplot as plt

import numpy as np

import seaborn as sns

import pandas as pd

import xarray as xr

sns.set(
    palette='husl',
    font_scale=2,
    font='serif',
)

fig_dir = 'results/figures'
os.makedirs(fig_dir, exist_ok=True)

# %% [code]
file_type_map = OrderedDict()
file_type_map['mz5'] = 'mz5 | msconvert | Profile'
file_type_map['mzML/a'] = 'mzML | msconvert | Profile'
file_type_map['mzML/g'] = 'mzML | Toffee Round Trip | Profile'
file_type_map['mzML/e'] = 'mzML | Docker | Profile'
file_type_map['mzML/b'] = 'mzML | Numpress | Profile'
file_type_map['mzML/f'] = 'mzML | Docker | Centroid'
file_type_map['mzML/c'] = 'mzML | msconvert | Centroid'
file_type_map['mzML/d'] = 'mzML | Numpress | Centroid'
file_type_map['tof/a'] = 'Toffee | msconvert | Profile'
file_type_map['tof/e'] = 'Toffee | Docker | Profile'
file_type_map['Vendor'] = 'Vendor'

order = [v for _, v in file_type_map.items()]

# set up colours
num_tof = 2
num_profile = 3
num_centroid = 3

colours = OrderedDict()
colours['mz5'] = plt.cm.Greens_r(0.1)
colours['mzML/a'] = plt.cm.Blues_r(0.1)
colours['mzML/g'] = plt.cm.Blues_r(0.2)
colours['mzML/e'] = plt.cm.Blues_r(0.3)
colours['mzML/b'] = plt.cm.Blues_r(0.4)
colours['mzML/f'] = plt.cm.Reds_r(0.1)
colours['mzML/c'] = plt.cm.Reds_r(0.2)
colours['mzML/d'] = plt.cm.Reds_r(0.3)
colours['tof/a'] = plt.cm.Purples_r(0.3)
colours['tof/e'] = plt.cm.Purples_r(0.4)
colours['Vendor'] = 'black'
for k, v in file_type_map.items():
    colours[v] = colours[k]

# %% [code]
def load_data(result_dir, ds_name, procan90=False):
    result = []
    for key, key_name in file_type_map.items():
        k = key.replace('/', '_')
        fname = f'{result_dir}/pqm_matrix-{k}.nc'
        if not os.path.exists(fname):
            continue
        ds = xr.open_dataset(fname)
        i = ds[ds_name].to_dataframe().reset_index()
        i['FileType'] = key
        i['File Type'] = key_name
        result.append(i)
    result = pd.concat(result)
    result['FDRIter'] = result.InjectionName.apply(lambda x: int(x.split('-')[-1]))
    if procan90:
        result['InjectionName'] = result.InjectionName.apply(lambda x: '-'.join(x.split('-')[:-1]))
    else:
        result['InjectionName'] = result.InjectionName.apply(lambda x: x.split('-')[0])
    return result

def create_confusion_matrix(rt_df, manual, nan_col, rt_threshold=10):
    # set up manual validation
    rt = pd.merge(rt_df, manual, on=['InjectionName', 'File Type', 'PQM'], how='outer')
    rt = rt[~rt[nan_col].isna()].reset_index(drop=True)  # drop those PQMs not found anywhere
    rt['Delta RT'] = rt['Expected Retention Time'] - rt['Retention Time']

    not_in_manual = rt['Expected Retention Time'].isna()
    not_in_run = rt['Retention Time'].isna()

    rt['True Negative'] = not_in_manual & not_in_run
    rt['False Negative'] = ~not_in_manual & not_in_run
    is_positive = ~rt['True Negative'] & ~rt['False Negative']

    rt['True Positive'] = (rt['Delta RT'].abs() <= rt_threshold) & is_positive
    rt['False Positive'] = ~rt['True Positive'] & is_positive

    rt['Total'] = 1  # for debugging
    
    return rt

def normalise_intensity_and_count_missing(
    intensity,
    metadata,
    normalising_sample,
    max_fraction_of_missing_per_pqm=1.0,
):
    index_cols = [
        'InjectionName',
        'FDRIter',
        'FileType',
        'PQM',
    ]
    i = intensity.set_index(index_cols).Intensity
    try:
        i = i.unstack(level=[2, 3])
    except:
        i = i.reset_index()
        m = i.duplicated(subset=index_cols, keep=False)
        display(i[m])
        raise

    # calculate missing
    is_missing = i.isna()
    frac_nans = is_missing.mean()
    nan_mask = frac_nans > max_fraction_of_missing_per_pqm

    is_missing.loc[:, nan_mask] = True
    is_missing = is_missing.stack().stack()
    is_missing.name = 'Missing'
    is_missing = is_missing.reset_index()
    
    # normalise intensity
    norm_vector = i.loc[pd.IndexSlice[normalising_sample, :], :].median()
    norm_i = i / norm_vector
    norm_i.loc[:, nan_mask] = np.NaN
    norm_i.fillna(0, inplace=True)  # fill nans with 0 so they're not lost in the reshape
    norm_i = norm_i.stack().stack().apply(np.log2)
    norm_i.name = 'Normalised Intensity'
    norm_i = norm_i.reset_index()
    
    # merge these two
    new_cols = pd.merge(norm_i, is_missing, on=index_cols, how='outer')
    assert new_cols.shape[0] == norm_i.shape[0], f'{new_cols.shape} != {norm_i.shape}'
    assert new_cols.shape[0] == is_missing.shape[0], f'{new_cols.shape} != {is_missing.shape}'

    # merge this with the intensity df
    result = pd.merge(intensity, new_cols, on=index_cols, how='outer')
    assert result.shape[0] >= intensity.shape[0], f'{intensity.shape} != {result.shape} != {new_cols.shape}'
    result = pd.merge(result, metadata, on='InjectionName')
    
    result.loc[result.Missing, 'Normalised Intensity'] = np.NaN
    return result

def gather_high_confidence_missing_pqms(
    norm_intensity,
    rt,
    qvalue,
    truth_file_type='mzML/e',
    other_file_type='mzML/f',
    number_to_collect=100,
):
    merge_cols = [
        'InjectionName',
        'FDRIter',
        'FileType',
        'File Type',
        'PQM',
    ]
    qvalue1 = pd.merge(
        norm_intensity[merge_cols + ['Missing', 'Intensity', 'Normalised Intensity']],
        qvalue[merge_cols + ['m_score']],
        on=merge_cols,
    )
    qvalue1 = pd.merge(
        qvalue1,
        rt[merge_cols + ['Retention Time']],
        on=merge_cols,
    )
    
    qvalue_e = qvalue1.loc[qvalue1.FileType == truth_file_type, :]
    qvalue_f = qvalue1.loc[qvalue1.FileType == other_file_type, :]
    merge_cols = [
        'InjectionName',
        'FDRIter',
        'PQM',
    ]
    qvalue2 = pd.merge(qvalue_e, qvalue_f, on=merge_cols, suffixes=['_true', '_other'])

    df = qvalue2[~qvalue2.Missing_true & qvalue2.Missing_other]
    merge_cols = ['InjectionName', 'PQM']
    
    count = df.groupby(merge_cols).FDRIter.count()
    name0 = 'NumberOfMissingFiles'
    count.name = name0
    count = count.to_frame()
    
    name1 = 'TotalQValue'
    count[name1] = df.groupby(merge_cols).m_score_true.sum()
    
    name2 = 'MedianQValue'
    count[name2] = df.groupby(merge_cols).m_score_true.median()
    
    name3 = 'AveQValueLog10Bin'
    count[name3] = count[name2].apply(lambda x: int(np.round(np.log10(x), 0)))

    count['RetentionTimeMedian'] = df.groupby(merge_cols)['Retention Time_true'].median()
    count['RetentionTimeMAD'] = df.groupby(merge_cols)['Retention Time_true'].mad()
    
    return count.sort_values([name3, name0, name2], ascending=[True, False, True]) \
        .head(n=number_to_collect) \
        .reset_index(drop=False)

# %% [markdown]
## File Sizes

# %% [code]
all_filesizes = []
files = {
    'SGS': 'results/filesizes/sgs-filesizes.tsv',
    'Roest2016': 'results/filesizes/roest2016-filesizes.tsv',
    'ProCan90': 'results/filesizes/procan90-filesizes.tsv',
}
for name, path in files.items():
    filesizes = pd.read_csv(path, sep='\t')
    # drop bad SGS file
    if name == 'SGS':
        filesizes = filesizes[~filesizes.InjectionName.str.contains('001')]
    filesizes.drop(['Scan', 'Wiff'], axis=1, inplace=True)
    filesizes.set_index('InjectionName', inplace=True)
    # filesizes /= filesizes.Vendor
    vendor = filesizes['Vendor'].copy()
    for c in filesizes.columns:
        filesizes[c] = filesizes[c] / vendor
    filesizes.columns.name = 'FileType'
    filesizes = filesizes.stack()
    filesizes.name = 'FileSize'
    filesizes = filesizes.reset_index()
    filesizes['Experiment'] = name
    all_filesizes.append(filesizes)

filesizes = pd.concat(all_filesizes)
filesizes['File Type'] = filesizes.FileType.map(file_type_map)

# %% [code]
fig, ax = plt.subplots(figsize=(20, 7))

ftypes = filesizes['File Type'].unique()
size_order = list(reversed([o for o in order if o in ftypes]))
palette = [colours[o] for o in size_order]

sns.barplot(
    data=filesizes,
    y='File Type',
    x='FileSize',
    palette=palette,
    ax=ax,
    linewidth=1,
    edgecolor='white',
    order=size_order,
    orient='h',
)
# ax.legend(loc=0, borderaxespad=1.0, frameon=False, prop={'size': 12})
ax.axvline(1, c='maroon', ls='--', lw=3, alpha=0.25)
_ = ax.set_xlabel('File size relative to vendor')
_ = ax.set_ylabel('')

fig.tight_layout()
fig.savefig(f'{fig_dir}/filesizes.pdf')
fig.savefig(f'{fig_dir}/filesizes.png')

# %% [markdown]
We can also extract data from the ProCan production systems to look at conversion times and runtimes. Note, for the pipeline run times, this is not a like-for-like comparison as things like the complexity of the spectral library and the command line arguments can be varied for different projects. Each of these changes will impact on the runtime of the pipeline.

# %% [code]
def load_prod_data():
    df = pd.read_csv('production-toffee-conversion-timings.csv.gz', low_memory=False)
    df_tof = df.loc[~df.TofUid.isna()] \
        .rename(columns={
            'TofUid': 'ArtifactUid',
            'TofFilesizeMB': 'ArtifactFilesizeMB',
            'TofPipelineWorkflowUid': 'PipelineWorkflowUid',
            'TofPipelineDate': 'PipelineDate',
            'TofPipelineRuntimeMinutes': 'PipelineRuntimeMinutes',
        }) \
        .dropna(how='all', axis=1)
    df_tof['ArtifactType'] = 'Toffee'

    df_mzml = df.loc[~df.MzmlUid.isna()] \
        .rename(columns={
            'MzmlUid': 'ArtifactUid',
            'MzmlFilesizeMB': 'ArtifactFilesizeMB',
            'MzmlPipelineWorkflowUid': 'PipelineWorkflowUid',
            'MzmlPipelineDate': 'PipelineDate',
            'MzmlPipelineRuntimeMinutes': 'PipelineRuntimeMinutes',
        }) \
        .dropna(how='all', axis=1)
    df_mzml['ArtifactType'] = 'MzML'
    df_mzml.loc[
        (df_mzml.ArtifactFilesizeMB / df_mzml.WiffFilesizeMB) < 2,
        'ArtifactType',
    ] = 'Centroid MzML'

    df = pd.concat((df_tof, df_mzml))
    bin_size = 400.0
    df['Wiff filesize (MB)'] = (
        bin_size * (0.5 + (df.WiffFilesizeMB / bin_size).astype(int))
    )
    
    return df

def plot_toffee_conversion_timing(production_filesizes):
    production_filesizes = production_filesizes.loc[
        (production_filesizes.ArtifactType == 'Toffee') &
        (production_filesizes.ConversionRuntimeMinutes > 0)
    ].copy()

    xname = 'Wiff filesize (MB)'
    yname = 'Wiff to Toffee (CPU hrs)'

    production_filesizes[yname] = production_filesizes.ConversionRuntimeMinutes / 60.0

    counts = production_filesizes[xname].groupby(production_filesizes[xname]).count()
    log_counts = counts.apply(np.log10)
    log_counts -= log_counts.min()
    log_counts /= log_counts.max()
    colours = log_counts.apply(lambda x: plt.cm.viridis(x, alpha=x)).tolist()

    fig, ax = plt.subplots(figsize=(20, 7))

    sns.barplot(
        data=production_filesizes,
        x=xname,
        y=yname,
        palette=colours,
        ax=ax,
        linewidth=1,
        edgecolor='white',
    )
    labels = []
    for i, l in enumerate(ax.xaxis.get_ticklabels()):
        labels.append(f'{float(l.get_text()):.0f}' if i % 2 == 0 else '')
    ax.set_xticklabels(labels)
    ax.set_ylabel(yname)
    ax.set_xlabel(xname)

    rects = [rect for rect in ax.get_children() if isinstance(rect, mpl.patches.Rectangle)]
    for rect, count in zip(rects, counts.tolist()):
        height = rect.get_height()
        if height < 40 / 60:
            yloc = 15
            clr = 'black'
            align = 'bottom'
        else:
            yloc = -15
            clr = 'white'
            align = 'top'
        ax.annotate(
            f'n = {count}',
            xy=(rect.get_x() + rect.get_width() / 2, height),
            xytext=(1, yloc),
            textcoords='offset points',
            ha='center',
            va=align,
            color=clr,
            weight='bold',
            fontsize=24,
            clip_on=True,
            rotation=90,
        )

    fig.tight_layout()
    fig.savefig(f'{fig_dir}/production-filesizes.pdf')
    fig.savefig(f'{fig_dir}/production-filesizes.png')
    plt.show()

def plot_workflow_timing(production_filesizes):
    production_filesizes = production_filesizes.loc[
        ~production_filesizes.PipelineWorkflowUid.isna()
    ].copy()

    xname = 'Wiff filesize (MB)'
    yname = 'Scoring pipeline (CPU hrs)'

    production_filesizes[yname] = production_filesizes.PipelineRuntimeMinutes / 60.0 * 15

    index = [
        (n, s)
        for n in production_filesizes['ArtifactType'].unique()
        for s in sorted(production_filesizes[xname].unique())
    ]
    counts = production_filesizes[xname].groupby([
        production_filesizes['ArtifactType'],
        production_filesizes[xname],
    ]).count().reindex(index)

    fig, ax = plt.subplots(figsize=(20, 7))

    sns.barplot(
        data=production_filesizes,
        x=xname,
        y=yname,
        hue='ArtifactType',
        palette='viridis',
        ax=ax,
        linewidth=1,
        edgecolor='white',
    )
    labels = []
    for i, l in enumerate(ax.xaxis.get_ticklabels()):
        labels.append(f'{float(l.get_text()):.0f}' if i % 2 == 0 else '')
    ax.set_xticklabels(labels)
    ax.set_ylabel(yname)
    ax.set_xlabel(xname)

    rects = [rect for rect in ax.get_children() if isinstance(rect, mpl.patches.Rectangle)]
    for rect, count in zip(rects, counts.tolist()):
        height = rect.get_height()
        if pd.isna(count):
            continue
        yloc = 30
        if height < 15 * 50 / 60:
            clr = 'black'
            align = 'bottom'
        else:
            yloc = -yloc
            clr = 'white'
            align = 'top'
        ax.annotate(
            f'n = {int(count)}',
            xy=(rect.get_x() + rect.get_width() / 2, height),
            xytext=(1, yloc),
            textcoords='offset points',
            ha='center',
            va=align,
            color=clr,
            weight='bold',
            fontsize=24,
            clip_on=True,
            rotation=90,
        )

    fig.tight_layout()
    fig.savefig(f'{fig_dir}/production-runtimes.pdf')
    fig.savefig(f'{fig_dir}/production-runtimes.png')
    plt.show()

production_filesizes = load_prod_data()
display(production_filesizes.head(n=3).T)
plot_toffee_conversion_timing(production_filesizes)
plot_workflow_timing(production_filesizes)

# %% [markdown]
## Swath Gold Standard

# %% [code]
sgs_result_dir = 'results/sgs'
sgs_rt = load_data(sgs_result_dir, ds_name='RT').rename(columns={'RT': 'Retention Time'})
sgs_intensity = load_data(sgs_result_dir, ds_name='Intensity')
sgs_qvalue = load_data(sgs_result_dir, ds_name='m_score')

# %% [code]
def load_metadata():
    metadata = pd.read_csv('sgs-metadata.tsv', sep='\t')
    mask1 = metadata.Background == 'Human'
    mask2 = metadata.TechnicalReplicate == 1
    metadata = metadata[mask1 & mask2]
    metadata = metadata.sort_values('Dilution', ascending=False).reset_index(drop=True)

    manual = pd.read_csv('manual_validation/OpenSWATH_SM3_GoldStandardManualResults.csv.gz')
    manual['PQM'] = manual.transition_group_id.apply(lambda x: x.split('_')[-1].replace('/', '_'))
    manual['InjectionName'] = manual.ReplicateName.str.lower()
    manual['RetentionTime'] *= 60
    manual = manual[manual.PeakRank == 1]
    cols = [
        'InjectionName',
        'PQM',
        'RetentionTime',
        'Area',
    ]
    manual = manual[manual.InjectionName.isin(metadata.InjectionName.unique())]
    manual = manual[cols].sort_values('InjectionName') \
        .reset_index(drop=True) \
        .rename(columns={'RetentionTime': 'Expected Retention Time'})
    manual = pd.merge(manual, metadata, on='InjectionName', how='outer')
    
    full_manual = []
    for k, v in file_type_map.items():
        if k == 'Vendor':
            continue
        manual['FileType'] = k
        manual['File Type'] = v
        full_manual.append(manual.copy())

    return metadata, pd.concat(full_manual)

sgs_metadata, sgs_manual = load_metadata()

# %% [markdown]
### Confusion Matrix

# %% [code]
sgs_confusion = create_confusion_matrix(sgs_rt, sgs_manual, nan_col='Dilution')
sgs_confusion['Dilution'] = sgs_confusion.Dilution.astype(int)
sgs_confusion['TechnicalReplicate'] = sgs_confusion.TechnicalReplicate.astype(int)
sgs_confusion = sgs_confusion[sgs_confusion.Dilution <= 32]

# %% [code]
cols = ['True Positive', 'False Positive', 'True Negative', 'False Negative', 'Total']
sgs_counts = sgs_confusion.groupby(['Dilution', 'InjectionName', 'FDRIter', 'File Type']).sum()[cols]
assert (
    sgs_counts[cols[0]] + 
    sgs_counts[cols[1]] + 
    sgs_counts[cols[2]] + 
    sgs_counts[cols[3]] == sgs_counts[cols[4]]
).all()
sgs_counts = sgs_counts.reset_index()

counts_order = [o for o in order if o in sgs_counts['File Type'].unique()]
palette = [colours[o] for o in counts_order]

# ---
fig, ax = plt.subplots(figsize=(15, 5))
sns.barplot(
    data=sgs_counts,
    x='Dilution',
    y='True Positive',
    hue='File Type',
    palette=palette,
    hue_order=counts_order,
    ax=ax,
)
ax.legend(bbox_to_anchor=(1.01, 0.5), loc=6, borderaxespad=0.0, frameon=False, prop={'size': 12})
fig.tight_layout()
fig.savefig(f'{fig_dir}/sgs-true-positive.pdf')
plt.show()

# ---
fig, ax = plt.subplots(figsize=(15, 5))
sns.barplot(
    data=sgs_counts,
    x='Dilution',
    y='False Positive',
    hue='File Type',
    palette=palette,
    hue_order=counts_order,
    ax=ax,
)
ax.legend(bbox_to_anchor=(1.01, 0.5), loc=6, borderaxespad=0.0, frameon=False, prop={'size': 12})
fig.tight_layout()
fig.savefig(f'{fig_dir}/sgs-false-positive.pdf')
plt.show()

# ---
fig, ax = plt.subplots(figsize=(15, 5))
sns.barplot(
    data=sgs_counts,
    x='Dilution',
    y='False Negative',
    hue='File Type',
    palette=palette,
    hue_order=counts_order,
    ax=ax,
)
ax.legend(bbox_to_anchor=(1.01, 0.5), loc=6, borderaxespad=0.0, frameon=False, prop={'size': 12})
fig.tight_layout()
fig.savefig(f'{fig_dir}/sgs-false-negative.pdf')
plt.show()

# %% [markdown]
### Intensities

# %% [code]
sgs_norm_intensity = normalise_intensity_and_count_missing(
    sgs_intensity,
    sgs_metadata,
    normalising_sample='napedro_l120417_010_sw',
    max_fraction_of_missing_per_pqm=1.0,
)

sgs_norm_intensity = sgs_norm_intensity[
    sgs_norm_intensity.Dilution <= 32
]

# %% [code]
df = gather_high_confidence_missing_pqms(
    sgs_norm_intensity,
    sgs_rt,
    sgs_qvalue,
    truth_file_type='mzML/e',
    other_file_type='mzML/a',
)
df.to_csv(f'{sgs_result_dir}/missing-profile-pqms.tsv', sep='\t', index=False)
df.head(n=2)

# %% [code]
df = gather_high_confidence_missing_pqms(
    sgs_norm_intensity,
    sgs_rt,
    sgs_qvalue,
    truth_file_type='mzML/e',
    other_file_type='mzML/f',
)
df.to_csv(f'{sgs_result_dir}/missing-centroid-pqms.tsv', sep='\t', index=False)
df.head(n=2)

# %% [code]
df = gather_high_confidence_missing_pqms(
    sgs_norm_intensity,
    sgs_rt,
    sgs_qvalue,
    truth_file_type='mzML/e',
    other_file_type='tof/e',
)
df.to_csv(f'{sgs_result_dir}/missing-toffee-pqms.tsv', sep='\t', index=False)
df.head(n=2)

# %% [code]
df = gather_high_confidence_missing_pqms(
    sgs_norm_intensity,
    sgs_rt,
    sgs_qvalue,
    truth_file_type='tof/e',
    other_file_type='mzML/e',
)
df.to_csv(f'{sgs_result_dir}/missing-mzml-pqms.tsv', sep='\t', index=False)
df.head(n=2)

# %% [code]
fig, ax = plt.subplots(figsize=(15, 6))

intensity_order = [o for o in order if o in sgs_intensity['File Type'].unique()]
palette = [colours[o] for o in intensity_order]

# ---
sns.boxplot(
    data=sgs_norm_intensity,
    x='Dilution',
    y='Normalised Intensity',
    hue='File Type',
    palette=palette,
    hue_order=intensity_order,
    ax=ax,
)
for x in ax.get_xticks():
    ax.plot(
        [x - 0.5, x + 0.5],
        [-x, -x],
        c='r',
        ls='--',
    )

ax.legend(bbox_to_anchor=(1.01, 0.5), loc=6, borderaxespad=0.0, frameon=False, prop={'size': 12})
fig.tight_layout()
fig.savefig(f'{fig_dir}/sgs-dilutions.pdf')
plt.show()

# %% [code]
fig, (ax1, ax2) = plt.subplots(nrows=2, figsize=(15, 7), sharex=True)

intensity_order = [o for o in order if o in sgs_intensity['File Type'].unique()]
palette = [colours[o] for o in intensity_order]

# ---
bias = sgs_norm_intensity['Normalised Intensity'].groupby([
    sgs_norm_intensity['File Type'],
    sgs_norm_intensity['Dilution'],
]).median().reset_index().rename(columns={'Normalised Intensity': 'Bias'})
bias['Bias'] = bias['Bias'] - (1 / bias.Dilution).apply(np.log2)

sns.stripplot(
    data=bias,
    x='Dilution',
    y='Bias',
    hue='File Type',
    palette=palette,
    hue_order=intensity_order,
    ax=ax1,
    dodge=True,
    s=25,
    linewidth=1,
)
ax1.legend(bbox_to_anchor=(1.01, 0.5), loc=6, borderaxespad=0.0, frameon=False, prop={'size': 12})
ax1.set_xlabel('')

#
variance = sgs_norm_intensity['Normalised Intensity'].groupby([
    sgs_norm_intensity['File Type'],
    sgs_norm_intensity['Dilution'],
]).mad().reset_index().rename(columns={'Normalised Intensity': 'Variance'})

sns.stripplot(
    data=variance,
    x='Dilution',
    y='Variance',
    hue='File Type',
    palette=palette,
    hue_order=intensity_order,
    ax=ax2,
    dodge=True,
    s=25,
    marker='v',
    linewidth=1,
)
ax2.legend_ = None

fig.tight_layout()
fig.savefig(f'{fig_dir}/sgs-bias-variance.pdf')
plt.show()

# %% [markdown]
## Roest2016

# %% [code]
roest2016_result_dir = 'results/roest2016'
roest2016_rt = load_data(roest2016_result_dir, ds_name='RT').rename(columns={'RT': 'Retention Time'})
roest2016_intensity = load_data(roest2016_result_dir, ds_name='Intensity')
roest2016_qvalue = load_data(roest2016_result_dir, ds_name='m_score')

# %% [code]
def load_metadata():
    metadata = pd.read_csv('roest2016-metadata.tsv', sep='\t')
    metadata = metadata.sort_values('InjectionName', ascending=False)

    manual = pd.read_csv('manual_validation/SkylineResult500Peptides.csv')
    manual['PQM'] = manual['Peptide Modified Sequence'] + '_' + manual['Precursor Charge'].map(str)
    manual['InjectionName'] = manual['Replicate Name']
    manual['Retention Time'] *= 60
    cols = [
        'InjectionName',
        'PQM',
        'Retention Time',
    ]
    manual = manual[manual.InjectionName.isin(metadata.InjectionName.unique())]
    manual = manual[cols].sort_values('InjectionName') \
        .reset_index(drop=True) \
        .rename(columns={'Retention Time': 'Expected Retention Time'})
    
    # filter for only those in the SRL
    srl = pd.read_csv('srl/roest2016/SpyogenesAssayLibrary.with_decoys.tsv', sep='\t', low_memory=False)
    srl = srl[srl.Decoy == 0].copy()
    srl['PQM'] = srl.ModifiedPeptideSequence + '_' + srl.PrecursorCharge.map(str)
    pqms = list(srl.PQM.unique())
    manual = manual[manual.PQM.isin(pqms)].copy()
    
    manual = pd.merge(manual, metadata, on='InjectionName', how='outer')
    
    full_manual = []
    for k, v in file_type_map.items():
        if k == 'Vendor':
            continue
        manual['FileType'] = k
        manual['File Type'] = v
        full_manual.append(manual.copy())

    return metadata, pd.concat(full_manual)

roest2016_metadata, roest2016_manual = load_metadata()

# %% [markdown]
### Confusion Matrix

# %% [code]
roest2016_confusion = create_confusion_matrix(roest2016_rt, roest2016_manual, nan_col='Condition')
roest2016_confusion['Biological Replicate'] = roest2016_confusion.BiologicalReplicate.astype(int)
roest2016_confusion['Technical Replicate'] = roest2016_confusion.TechnicalReplicate.astype(int)

# %% [code]
cols = ['True Positive', 'False Positive', 'True Negative', 'False Negative', 'Total']
roest2016_counts = roest2016_confusion.groupby([
    'Condition',
    'InjectionName',
    'FDRIter',
    'File Type',
    'Biological Replicate',
    'Technical Replicate',
]).sum()[cols]
assert (
    roest2016_counts[cols[0]] + 
    roest2016_counts[cols[1]] + 
    roest2016_counts[cols[2]] + 
    roest2016_counts[cols[3]] == roest2016_counts[cols[4]]
).all()
roest2016_counts = roest2016_counts.reset_index()

counts_order = [o for o in order if o in roest2016_counts['File Type'].unique()]
palette = [colours[o] for o in counts_order]

# ---
fig, ax = plt.subplots(figsize=(15, 5))
sns.barplot(
    data=roest2016_counts,
    x='Condition',
    y='True Positive',
    hue='File Type',
    palette=palette,
    hue_order=counts_order,
    ax=ax,
)
ax.legend(bbox_to_anchor=(1.01, 0.5), loc=6, borderaxespad=0.0, frameon=False, prop={'size': 12})
fig.tight_layout()
fig.savefig(f'{fig_dir}/roest2016-true-positive.pdf')
plt.show()

# ---
fig, ax = plt.subplots(figsize=(15, 5))
sns.barplot(
    data=roest2016_counts,
    x='Condition',
    y='False Positive',
    hue='File Type',
    palette=palette,
    hue_order=counts_order,
    ax=ax,
)
ax.legend(bbox_to_anchor=(1.01, 0.5), loc=6, borderaxespad=0.0, frameon=False, prop={'size': 12})
fig.tight_layout()
fig.savefig(f'{fig_dir}/roest2016-false-positive.pdf')
plt.show()

# ---
fig, ax = plt.subplots(figsize=(15, 5))
sns.barplot(
    data=roest2016_counts,
    x='Condition',
    y='False Negative',
    hue='File Type',
    palette=palette,
    hue_order=counts_order,
    ax=ax,
)
ax.legend(bbox_to_anchor=(1.01, 0.5), loc=6, borderaxespad=0.0, frameon=False, prop={'size': 12})
fig.tight_layout()
fig.savefig(f'{fig_dir}/roest2016-false-negative.pdf')
plt.show()

# %% [markdown]
### Intensities

# %% [code]
roest2016_norm_intensity = normalise_intensity_and_count_missing(
    roest2016_intensity,
    roest2016_metadata,
    normalising_sample='Strep0Repl1_R01_SW',
    max_fraction_of_missing_per_pqm=1.0,
)
roest2016_norm_intensity['Biological Replicate'] = roest2016_norm_intensity.BiologicalReplicate.astype(int)
roest2016_norm_intensity['Technical Replicate'] = roest2016_norm_intensity.TechnicalReplicate.astype(int)

# %% [code]
df = gather_high_confidence_missing_pqms(
    roest2016_norm_intensity,
    roest2016_rt,
    roest2016_qvalue,
    truth_file_type='mzML/e',
    other_file_type='mzML/a',
)
df.to_csv(f'{roest2016_result_dir}/missing-profile-pqms.tsv', sep='\t', index=False)
df.head(n=2)

# %% [code]
df = gather_high_confidence_missing_pqms(
    roest2016_norm_intensity,
    roest2016_rt,
    roest2016_qvalue,
    truth_file_type='mzML/e',
    other_file_type='mzML/f',
)
df.to_csv(f'{roest2016_result_dir}/missing-centroid-pqms.tsv', sep='\t', index=False)
df.head(n=2)

# %% [code]
df = gather_high_confidence_missing_pqms(
    roest2016_norm_intensity,
    roest2016_rt,
    roest2016_qvalue,
    truth_file_type='mzML/e',
    other_file_type='tof/e',
)
df.to_csv(f'{roest2016_result_dir}/missing-toffee-pqms.tsv', sep='\t', index=False)
df.head(n=2)

# %% [code]
df = gather_high_confidence_missing_pqms(
    roest2016_norm_intensity,
    roest2016_rt,
    roest2016_qvalue,
    truth_file_type='tof/e',
    other_file_type='mzML/e',
)
df.to_csv(f'{roest2016_result_dir}/missing-mzml-pqms.tsv', sep='\t', index=False)
df.head(n=2)

# %% [code]
num_pqms = roest2016_norm_intensity.PQM.unique().size

roest2016_missing_counts = roest2016_norm_intensity.set_index([
    roest2016_norm_intensity['InjectionName'],
    roest2016_norm_intensity['FDRIter'],
    roest2016_norm_intensity['Condition'],
    roest2016_norm_intensity['Biological Replicate'],
    roest2016_norm_intensity['Technical Replicate'],
    roest2016_norm_intensity['FileType'],
    roest2016_norm_intensity['File Type'],
    roest2016_norm_intensity['PQM'],
]).Missing.unstack().fillna(value=True).sum(axis=1)

as_percent = True
if as_percent:
    roest2016_missing_counts /= num_pqms
    roest2016_missing_counts *= 100
    name = 'Missing Values (%)'
else:
    name = 'Missing Values'

roest2016_missing_counts.name = name
roest2016_missing_counts = roest2016_missing_counts.reset_index()

missing_counts_order = [o for o in order if o in roest2016_missing_counts['File Type'].unique()]
palette = [colours[o] for o in missing_counts_order]

# ---
fig, ax = plt.subplots(figsize=(15, 5))
ax.set_title(f'Number of PQMs = {num_pqms}')
sns.barplot(
    data=roest2016_missing_counts,
    x='Condition',
    y=name,
    hue='File Type',
    palette=palette,
    hue_order=missing_counts_order,
    ax=ax,
)
ax.legend(bbox_to_anchor=(1.01, 0.5), loc=6, borderaxespad=0.0, frameon=False, prop={'size': 12})
fig.tight_layout()
fig.savefig(f'{fig_dir}/roest2016-missing-values.pdf')
plt.show()

# %% [markdown]
## ProCan90

# %% [code]
procan90_result_dir = 'results/procan90'
procan90_rt = load_data(procan90_result_dir, ds_name='RT', procan90=True).rename(columns={'RT': 'Retention Time'})
procan90_intensity = load_data(procan90_result_dir, ds_name='Intensity', procan90=True)
procan90_qvalue = load_data(procan90_result_dir, ds_name='m_score', procan90=True)
procan90_metadata = pd.DataFrame(
    [{'InjectionName': 'ProCan90-M0{}-01'.format(m), 'Machine': m} for m in range(1, 7)],
)

# %% [markdown]
### Intensities

# %% [code]
procan90_norm_intensity = normalise_intensity_and_count_missing(
    procan90_intensity,
    procan90_metadata,
    normalising_sample='ProCan90-M01-01',
    max_fraction_of_missing_per_pqm=1.0,
)

# %% [code]
df = gather_high_confidence_missing_pqms(
    procan90_norm_intensity,
    procan90_rt,
    procan90_qvalue,
    truth_file_type='mzML/e',
    other_file_type='mzML/a',
)
df.to_csv(f'{procan90_result_dir}/missing-profile-pqms.tsv', sep='\t', index=False)
df.head(n=2)

# %% [code]
df = gather_high_confidence_missing_pqms(
    procan90_norm_intensity,
    procan90_rt,
    procan90_qvalue,
    truth_file_type='mzML/e',
    other_file_type='mzML/f',
)
df.to_csv(f'{procan90_result_dir}/missing-centroid-pqms.tsv', sep='\t', index=False)
df.head(n=2)

# %% [code]
df = gather_high_confidence_missing_pqms(
    procan90_norm_intensity,
    procan90_rt,
    procan90_qvalue,
    truth_file_type='mzML/e',
    other_file_type='tof/e',
)
df.to_csv(f'{procan90_result_dir}/missing-toffee-pqms.tsv', sep='\t', index=False)
df.head(n=2)

# %% [code]
df = gather_high_confidence_missing_pqms(
    procan90_norm_intensity,
    procan90_rt,
    procan90_qvalue,
    truth_file_type='tof/e',
    other_file_type='mzML/e',
)
df.to_csv(f'{procan90_result_dir}/missing-mzml-pqms.tsv', sep='\t', index=False)
df.head(n=2)

# %% [code]
num_pqms = procan90_norm_intensity.PQM.unique().size

procan90_missing_counts = procan90_norm_intensity.set_index([
    procan90_norm_intensity['InjectionName'],
    procan90_norm_intensity['FDRIter'],
    procan90_norm_intensity['Machine'],
    procan90_norm_intensity['FileType'],
    procan90_norm_intensity['File Type'],
    procan90_norm_intensity['PQM'],
]).Missing.unstack().fillna(value=True).sum(axis=1)

as_percent = True
if as_percent:
    procan90_missing_counts /= num_pqms
    procan90_missing_counts *= 100
    name = 'Missing Values (%)'
else:
    name = 'Missing Values'

procan90_missing_counts.name = name
procan90_missing_counts = procan90_missing_counts.reset_index()

missing_counts_order = [o for o in order if o in procan90_missing_counts['File Type'].unique()]
palette = [colours[o] for o in missing_counts_order]

fig, ax = plt.subplots(figsize=(15, 5))
ax.set_title(f'Number of PQMs = {num_pqms}')
g = sns.barplot(
    data=procan90_missing_counts,
    x='Machine',
    y=name,
    hue='File Type',
    palette=palette,
    hue_order=missing_counts_order,
    ax=ax,
)
ax.legend(bbox_to_anchor=(1.01, 0.5), loc=6, borderaxespad=0.0, frameon=False, prop={'size': 12})
fig.tight_layout()
fig.savefig(f'{fig_dir}/procan90-missing-values.pdf')
plt.show()

# %% [code]
fig, ax = plt.subplots(figsize=(15, 5))

intensity_order = [o for o in order if o in procan90_intensity['File Type'].unique()]
palette = [colours[o] for o in intensity_order]

# ---
sns.boxplot(
    data=procan90_norm_intensity,
    x='Machine',
    y='Normalised Intensity',
    hue='File Type',
    palette=palette,
    hue_order=intensity_order,
    ax=ax,
)
for x in ax.get_xticks():
    ax.plot(
        [x - 0.5, x + 0.5],
        [0, 0],
        c='r',
        ls='--',
    )

ax.legend(bbox_to_anchor=(1.01, 0.5), loc=6, borderaxespad=0.0, frameon=False, prop={'size': 12})
fig.tight_layout()
fig.savefig(f'{fig_dir}/procan90-dilution-series.pdf')
plt.show()

# %% [markdown]
## Combined figures

# %% [code]
import matplotlib.gridspec as gridspec

fig, ax = plt.subplots(figsize=(20, 13))

gs = gridspec.GridSpec(3, 4)
gs.update(wspace=0.55, hspace=0.55)

roest2016_tp = plt.subplot(gs[1, 0])
roest2016_fp = plt.subplot(gs[1, 1])
roest2016_fn = plt.subplot(gs[1, 2], sharey=roest2016_fp)
roest2016_missing = plt.subplot(gs[2, 0])
procan90_missing = plt.subplot(gs[2, 1:], sharey=roest2016_missing)
sgs_tp = plt.subplot(gs[0, :2])
sgs_dilution = plt.subplot(gs[0, 2:])

kwargs = dict(
    xy=(-50, 200),
    xycoords='axes points',
    va='center',
    ha='left',
    fontsize=20,
    weight='bold',
)
label_fontsize = 16

sgs_tp.annotate('A.', **kwargs)
sgs_dilution.annotate('B.', **kwargs)
roest2016_tp.annotate('C.', **kwargs)
roest2016_missing.annotate('D.', **kwargs)

# ******************
# ---
hue_order = [o for o in order if o in sgs_counts['File Type'].unique()]
palette = [colours[o] for o in hue_order]
ax = sgs_tp
sns.barplot(
    data=sgs_counts,
    x='Dilution',
    y='True Positive',
    hue='File Type',
    palette=palette,
    hue_order=hue_order,
    ax=ax,
)
ax.legend_ = None
ax.set_title(ax.yaxis.get_label().get_text())
ax.set_ylabel('Count', fontsize=label_fontsize)
ax.set_xlabel('Dilution', fontsize=label_fontsize)

# ******************
hue_order = [o for o in order if o in roest2016_counts['File Type'].unique()]
palette = [colours[o] for o in hue_order]
ax = roest2016_tp
sns.barplot(
    data=roest2016_counts,
    x='Condition',
    y='True Positive',
    hue='File Type',
    palette=palette,
    hue_order=hue_order,
    ax=ax,
)
ax.legend_ = None
ax.set_title(ax.yaxis.get_label().get_text())
ax.set_ylabel('Count', fontsize=label_fontsize)
ax.set_xlabel('Condition', fontsize=label_fontsize)

ax = roest2016_fp
sns.barplot(
    data=roest2016_counts,
    x='Condition',
    y='False Positive',
    hue='File Type',
    palette=palette,
    hue_order=hue_order,
    ax=ax,
)
ax.legend_ = None
ax.set_title(ax.yaxis.get_label().get_text())
ax.set_ylabel('Count', fontsize=label_fontsize)
ax.set_xlabel('Condition', fontsize=label_fontsize)

ax = roest2016_fn
sns.barplot(
    data=roest2016_counts,
    x='Condition',
    y='False Negative',
    hue='File Type',
    palette=palette,
    hue_order=hue_order,
    ax=ax,
)
ax.legend(bbox_to_anchor=(1.1, 0.5), loc=6, borderaxespad=0.0, frameon=False, prop={'size': 14})
ax.set_title(ax.yaxis.get_label().get_text())
ax.set_ylabel('Count', fontsize=label_fontsize)
ax.set_xlabel('Condition', fontsize=label_fontsize)

# ******************
# roest2016_missing.set_title(f'Number of PQMs = {num_pqms}')
hue_order = [o for o in order if o in roest2016_missing_counts['File Type'].unique()]
palette = [colours[o] for o in hue_order]
ax = roest2016_missing
sns.barplot(
    data=roest2016_missing_counts,
    x='Condition',
    y=name,
    hue='File Type',
    palette=palette,
    hue_order=hue_order,
    ax=ax,
)
ax.legend_ = None
ax.set_title(ax.yaxis.get_label().get_text())
ax.set_ylabel('Percent', fontsize=label_fontsize)
ax.set_xlabel('Condition', fontsize=label_fontsize)

hue_order = [o for o in order if o in procan90_missing_counts['File Type'].unique()]
palette = [colours[o] for o in hue_order]
ax = procan90_missing
g = sns.barplot(
    data=procan90_missing_counts,
    x='Machine',
    y=name,
    hue='File Type',
    palette=palette,
    hue_order=hue_order,
    ax=ax,
)
ax.legend_ = None
ax.set_title(ax.yaxis.get_label().get_text())
ax.set_ylabel('Percent', fontsize=label_fontsize)
ax.set_xlabel('Mass Spectrometer', fontsize=label_fontsize)

# ******************
hue_order = [o for o in order if o in sgs_norm_intensity['File Type'].unique()]
palette = [colours[o] for o in hue_order]
ax = sgs_dilution
sns.boxplot(
    data=sgs_norm_intensity,
    x='Dilution',
    y='Normalised Intensity',
    hue='File Type',
    palette=palette,
    hue_order=hue_order,
    ax=ax,
)
for x in ax.get_xticks():
    ax.plot(
        [x - 0.5, x + 0.5],
        [-x, -x],
        c='r',
        ls='--',
    )
ax.legend_ = None
ax.set_title(ax.yaxis.get_label().get_text())
ax.set_ylabel('Log2', fontsize=label_fontsize)
ax.set_xlabel('Dilution', fontsize=label_fontsize)

# fig.tight_layout()
fig.savefig(f'{fig_dir}/confusion-matrix.pdf')
plt.show()

# %% [code]
```
