## Supplementary material for "Toffee – a highly efficient, lossless file format for DIA-MS": Jupyter notebooks: calculate_ims.ipynb

```
# %% [code]
import h5py
from enum import Enum

import matplotlib.pyplot as plt

import numpy as np

import pandas as pd

import scipy.optimize

import seaborn as sns

from tqdm import tqdm

sns.set(
    palette='husl',
    font_scale=2,
)

# %% [code]
class IMSType(Enum):
    TOF = 'TOF'
    ORBITRAP = 'Orbitrap'


def load_dataset(spec_fname, ims_type=IMSType.TOF):
    if ims_type == IMSType.TOF:
        transfer_func = np.sqrt
    elif ims_type == IMSType.ORBITRAP:
        transfer_func = lambda x: 1.0 / np.sqrt(x)
    else:
        raise ValueError(f'Invalid IMS type: {ims_type}')

    result = {}
    with h5py.File(spec_fname, 'r') as f:
        for window in f.keys():
            print(window, flush=True)
            mz = []
            transformed_mz = []
            delta_transformed_mz = []
            window_indices = [0]
            rt = []
            for scan in tqdm(f[window].keys()):
                s = f[window][scan]
                tmp_mz = s['MASS_OVER_CHARGE'][()]
                
                rt.append(s.attrs['RETENTION_TIME'])
                
                n_points = tmp_mz.shape[0]
                window_indices.append(n_points)
                if n_points == 0:
                    continue
                
                tmp_transformed_mz = transfer_func(tmp_mz)
                tmp_delta_transformed_mz = [tmp_transformed_mz[0]]
                if tmp_transformed_mz.shape[0] > 1:
                    tmp_delta_transformed_mz += list(np.diff(tmp_transformed_mz))
                
                assert tmp_transformed_mz.shape[0] == n_points
                assert len(tmp_delta_transformed_mz) == n_points
                mz.extend(tmp_mz)
                transformed_mz.extend(tmp_transformed_mz)
                delta_transformed_mz.extend(tmp_delta_transformed_mz)
            
            indices = np.cumsum(window_indices)
            df = pd.DataFrame(
                {
                    'MZ': mz,
                    'TransformedMZ': transformed_mz,
                    'DeltaTransformedMZ': delta_transformed_mz,
                },
            )
            result[window] = {
                'Indices': indices,
                'RetentionTime': rt,
                'MZDataFrame': df,
            }
        
    return result


def calculate_alpha_and_beta_directly(window_name, window_data, ims_type=IMSType.TOF, output_basename=None):

    def calc_mz(idx, alpha, beta):
        transformed = alpha * idx + beta
        if ims_type == IMSType.TOF:
            return transformed ** 2
        elif ims_type == IMSType.ORBITRAP:
            return transformed ** -2
        else:
            raise ValueError(f'Invalid IMS type: {ims_type}')

    def guess_alpha(transformed):
        if ims_type == IMSType.TOF:
            return np.min(transformed)
        elif ims_type == IMSType.ORBITRAP:
            return np.max(transformed)
        else:
            raise ValueError(f'Invalid IMS type: {ims_type}')
    
    indices = window_data['Indices']
    rt_vec = window_data['RetentionTime']
    df = window_data['MZDataFrame']

    mz = 1.0 * df.MZ.values
    r = 1.0 * df.TransformedMZ.values
    dr = 1.0 * df.DeltaTransformedMZ.values
    rt_full_vec = np.zeros_like(mz)
    for i, rt in enumerate(rt_vec):
        start = indices[i]
        end = indices[i + 1]
        dr[start] = 0.0
        rt_full_vec[start:end] = rt

    # ---
    # calculate alpha
    mask = dr != 0
    alpha_start = guess_alpha(dr[mask])
    alpha = alpha_start
    di = np.round(dr / alpha).astype(int)
    alpha_full_vec = np.zeros_like(mz)
    alpha_full_vec[~mask] = np.NaN
    for i in range(10):
        alpha_full_vec[mask] = dr[mask] / di[mask]
        alpha = np.nanmedian(alpha_full_vec)
        di_new = np.round(dr / alpha).astype(int)
        delta_di = np.sum(np.abs(di - di_new))
        di = di_new
        if delta_di == 0:
            break
        assert i != 9
    assert np.min(di[mask]) >= 1, 'Delta Index cannot have values less than 1, it has: ' + str(np.sum(di[mask] < 1))
    
    fig, ax = plt.subplots(figsize=(15, 5))
    if ims_type == IMSType.TOF:
        mask1 = mask & (dr < 20 * alpha)
    elif ims_type == IMSType.ORBITRAP:
        mask1 = mask & (dr > 20 * alpha)
    else:
        raise ValueError(f'Invalid IMS type: {ims_type}')
    _mz, _dr = mz[mask1], dr[mask1]
    mask1 = np.random.choice(_mz.shape[0], min(10000, _mz.shape[0]))
    ax.plot(
        _mz[mask1],
        _dr[mask1],
        'k.',
    )
    ax.axhline(alpha, ls='--', label='Multiples of Alpha')
    for i in range(2, 21):
        ax.axhline(i * alpha, ls='--')
    ax.legend(loc='upper right')
    ax.set_ylabel('Delta Transformed m/z')
    ax.set_xlabel('m/z')
    ax.set_title(window_name.upper())
    plt.show()

    # ---
    # calculate beta
    idx_full_vec = np.zeros_like(mz)
    beta_full_vec = np.zeros_like(mz)
    ppm_full_vec = np.zeros_like(mz)
    _idx0_vec = []
    _alpha_vec = []
    _beta_vec = []
    for i, rt in enumerate(rt_vec):
        start = indices[i]
        end = indices[i + 1]
        idx_vec = np.cumsum(di[start:end])

        alpha = np.nanmedian(alpha_full_vec[start:end])
        beta_vec = r[start:end] - alpha * idx_vec
        beta = np.nanmedian(beta_vec)

        idx0 = np.round(beta / alpha).astype(int)
        beta_vec -= idx0 * alpha
        beta -= idx0 * alpha
        
        if beta > 0:
            idx0 += 1
            beta_vec -= alpha
            beta -= alpha

        idx_full_vec[start:end] = idx0 + idx_vec
        beta_full_vec[start:end] = beta_vec
        _idx0_vec.append(idx0)
        _alpha_vec.append(alpha)
        _beta_vec.append(beta)
        
        new_mz = calc_mz(idx_full_vec[start:end], alpha, beta)
        ppm = 1e6 * (new_mz - mz[start:end]) / mz[start:end]
        ppm_full_vec[start:end] = ppm

    idx0_vec = np.array(_idx0_vec)
    alpha_vec = np.array(_alpha_vec)
    beta_vec = np.array(_beta_vec)
        
    alpha = np.nanmedian(alpha_full_vec)
    beta = np.nanmedian(beta_full_vec)

    ppm_median_full_vec = 1e6 * (calc_mz(idx_full_vec, alpha, beta) - mz) / mz

    fig, (ax_alpha, ax_beta) = plt.subplots(nrows=2, sharex=True, figsize=(20, 10))
    
    ax_alpha.plot(rt_vec, alpha_vec, 'k.')
    ax_alpha.axhline(alpha, ls='--', label=f'Alpha rel. to {alpha:.5e}')
    ax_alpha.set_ylabel(f'Alpha')
    ax_alpha.legend(loc='upper right')
    ax_alpha.set_title(window_name.upper())

    ax_beta.plot(rt_vec, beta_vec, 'k.')
    ax_beta.axhline(beta, ls='--', label=f'Beta rel. to {beta:.5e}')
    ylim = np.percentile(beta_vec, [5, 95])
    ylim_delta = ylim[1] - ylim[0]
    ylim_ave = 0.5 * (ylim[0] + ylim[1])
    ylim = [ylim_ave - ylim_delta, ylim_ave + ylim_delta]
    ax_beta.set_ylim(ylim)
    ax_beta.set_ylabel(f'Beta')
    ax_beta.legend(loc='upper right')
    ax_beta.set_xlabel('Retention Time (s)')

    fig.tight_layout()
    plt.show()
    
    fig, (ax1, ax2) = plt.subplots(nrows=2, sharex=True, figsize=(20, 10))
    np.random.seed(0)
    mask = np.random.choice(mz.shape[0], min(50000, mz.shape[0]))
    ax1.scatter(
        x=mz[mask],
        y=ppm_full_vec[mask],
        s=4,
        c=rt_full_vec[mask],
        alpha=0.25,
        cmap=plt.cm.viridis,
    )
    ylim = np.percentile(ppm_full_vec, [1, 99])
    ylim_delta = ylim[1] - ylim[0]
    ylim_ave = 0.5 * (ylim[0] + ylim[1])
    ylim = [ylim_ave - ylim_delta, ylim_ave + ylim_delta]
    ax1.set_ylim(ylim)
    ax1.set_ylabel('PPM (scan)')
    ax1.set_title(window_name.upper())
    
    ax2.scatter(
        x=mz[mask],
        y=ppm_median_full_vec[mask],
        s=4,
        c=rt_full_vec[mask],
        alpha=0.25,
        cmap=plt.cm.viridis,
    )
    ylim = np.percentile(ppm_median_full_vec, [1, 99])
    ylim_delta = ylim[1] - ylim[0]
    ylim_ave = 0.5 * (ylim[0] + ylim[1])
    ylim = [ylim_ave - ylim_delta, ylim_ave + ylim_delta]
    ax2.set_ylim(ylim)
    ax2.set_ylabel('PPM (window)')
    ax2.set_xlabel('m/z')
    
    fig.tight_layout()
    plt.show()
    
    # save for testing
    if output_basename is not None:
        gamma = idx_full_vec.min() - 10
        idx_full_vec -= gamma
        indices = indices[1:]
        assert alpha_vec.shape[0] == indices.shape[0]
        assert beta_vec.shape[0] == indices.shape[0]
        with h5py.File(output_basename, 'w') as f:
            f.attrs['median_alpha'] = alpha
            f.attrs['median_beta'] = beta
            f.attrs['gamma'] = int(gamma)
            def add_dataset(name, data):
                f.create_dataset(name, data=data, compression='gzip', compression_opts=6, chunks=data.shape)
            add_dataset('mz', mz)
            add_dataset('alpha', alpha_vec)
            add_dataset('beta', beta_vec)
            add_dataset('coord', idx_full_vec.astype(np.uint32))
            add_dataset('slice_index', indices.astype(np.uint32))

# %% [markdown]
## TripleTOF 6600

# %% [markdown]
### Water only injection #1

# %% [code]
blank_1 = load_dataset('raw_data/blank/180508_blank_S_M02_1.e.ms2-050.spech5')

# %% [code]
for window, window_data in blank_1.items():
    if window.startswith('ms2'):
        output_basename = f'raw_data/blank/180508_blank_S_M02_1.e.IMSProps.{window}.h5'
    else:
        output_basename = None
    calculate_alpha_and_beta_directly(window, window_data, output_basename=output_basename)

# %% [markdown]
### Water only injection #2

# %% [code]
blank_2 = load_dataset('raw_data/blank/180508_blank_S_M02_2.e.ms2-050.spech5')

# %% [code]
for window, window_data in blank_2.items():
    if window.startswith('ms2'):
        output_basename = f'raw_data/blank/180508_blank_S_M02_2.e.IMSProps.{window}.h5'
    else:
        output_basename = None
    calculate_alpha_and_beta_directly(window, window_data, output_basename=output_basename)

# %% [markdown]
### ProCan90-M03-01 | msconvert

# %% [code]
procan90_a = load_dataset('raw_data/procan90/ProCan90-M03-01.a.ms2-050.spech5')

# %% [code]
for window, window_data in procan90_a.items():
    calculate_alpha_and_beta_directly(window, window_data)

# %% [markdown]
### ProCan90-M03-01 | msconvert + numpress

# %% [code]
procan90_b = load_dataset('raw_data/procan90/ProCan90-M03-01.b.ms2-050.spech5')

# %% [code]
for window, window_data in procan90_b.items():
    calculate_alpha_and_beta_directly(window, window_data)

# %% [markdown]
### ProCan90-M03-01 | Sciex Docker

# %% [code]
procan90_e = load_dataset('raw_data/procan90/ProCan90-M03-01.e.ms2-050.spech5')

# %% [code]
for window, window_data in procan90_e.items():
    calculate_alpha_and_beta_directly(window, window_data)

# %% [markdown]
## TripleTOF 5600

# %% [code]
sgs = load_dataset('raw_data/sgs/napedro_l120227_004_sw.ms2-016.spech5')

# %% [code]
for window, window_data in sgs.items():
    if window.startswith('ms2'):
        output_basename = f'raw_data/sgs/napedro_l120227_004_sw.IMSProps.{window}.h5'
    else:
        output_basename = None
    calculate_alpha_and_beta_directly(window, window_data, output_basename=output_basename)

# %% [code]
```
