## Supplementary material for "Toffee – a highly efficient, lossless file format for DIA-MS": Jupyter notebooks: in-silico-dilution-plot.ipynb

```
# %% [markdown]
# In-silico Dilution of Toffee Data

With the advantages of `toffee` we can create an in-silico dilution data set for further analysis of pipelines, such as global FDR control, where we have a ground truth.

# %% [code]
%load_ext autoreload
%autoreload 2

# %% [code]
import os

import matplotlib.gridspec as gridspec
import matplotlib.pyplot as plt

import numpy as np

import seaborn as sns

import pandas as pd

import plotly.io as pio

import toffee
from toffee.util import calculate_isotope_mz
from toffee.viz import ToffeeFragmentsPlotter

base_dir = os.environ.get('DIA_TEST_DATA_REPO', None)
assert base_dir is not None
fig_ext = 'svg'

sns.set(
    palette='husl',
    font_scale=2,
    font='serif',
)

tof_dir = f'{base_dir}/in-silico-dilution/tof'
base_output_dir = 'results/in-silico-dilution'
results_path = f'{base_output_dir}/raw'
fig_dir = 'results/figures'

# %% [code]
toffee.__version__

# %% [markdown]
Load the spectral library

# %% [code]
def load_srl(srl_path):
    srl = pd.read_csv(srl_path, sep='\t')
    srl['PrecursorId'] = srl.ModifiedPeptideSequence + '_' + srl.PrecursorCharge.map(str)
    srl = srl[srl.Decoy == 0]
    return srl

srl = load_srl(f'{base_dir}/in-silico-dilution/srl/hek-top2000.with_decoys.tsv')

# pqm = 'AAFTAFEEAQLPR_2'
pqm = 'SKPGAAMVEMADGYAVDR_3'  # top_results.peptide_group_label.iloc[1]

n_isotopes = 0
precursor, product = calculate_isotope_mz(
    precursor_and_product_df=srl,
    n_isotopes=n_isotopes,
    transition_group_id=pqm,
    sort_by_intensity=True,
)

# %% [markdown]
Set up the plotters

# %% [code]
lower_mz_overlap = 0.0
upper_mz_overlap = 1.0

foreground_plotter = ToffeeFragmentsPlotter(
    f'{tof_dir}/ProCan90-M06-07.tof',
    lower_mz_overlap=lower_mz_overlap,
    upper_mz_overlap=upper_mz_overlap,
)

background_plotter = ToffeeFragmentsPlotter(
    f'{tof_dir}/190109_Missing-1-Bcontrol-RT_007V1_007WJ_M05_S_1.tof',
    lower_mz_overlap=lower_mz_overlap,
    upper_mz_overlap=upper_mz_overlap,
)

dilution = 1
combined_plotter = ToffeeFragmentsPlotter(
    f'{tof_dir}/in-silico-dilution.procan90-m06-07.ecoli.{dilution:02d}.tof',
    lower_mz_overlap=lower_mz_overlap,
    upper_mz_overlap=upper_mz_overlap,
)

# %% [markdown]
Plot the foreground, background and combination

# %% [code]
fig, ax = plt.subplots(figsize=(20, 13))

gs = gridspec.GridSpec(
    ncols=2,
    nrows=4,
    figure=fig,
    width_ratios=(4, 1),
    wspace=0.01,
    hspace=0.01,
)

bg = plt.subplot(gs[1, 0])
fg = plt.subplot(gs[0, 0], sharex=bg)
cb = plt.subplot(gs[2, 0], sharex=bg)
chrom = plt.subplot(gs[3, 0], sharex=bg)

bg_zoom = plt.subplot(gs[1, 1])
fg_zoom = plt.subplot(gs[0, 1], sharex=bg_zoom)
cb_zoom = plt.subplot(gs[2, 1], sharex=bg_zoom)
chrom_zoom = plt.subplot(gs[3, 1], sharex=bg_zoom)

retention_time_zoom = (2300.0, 2750.0)
details = [
    ('Foreground', fg_zoom, foreground_plotter, retention_time_zoom, 2, plt.cm.Dark2(0)),
    ('Background', bg_zoom, background_plotter, retention_time_zoom, 1, plt.cm.Dark2(7)),
    ('Combined', cb_zoom, combined_plotter, retention_time_zoom, 3, plt.cm.Dark2(3)),
    ('Foreground', fg, foreground_plotter, None, 2, plt.cm.Dark2(0)),
    ('Background', bg, background_plotter, None, 1, plt.cm.Dark2(7)),
    ('Combined', cb, combined_plotter, None, 3, plt.cm.Dark2(3)),
]
normalise_per_fragment = False

pqp_precursors = precursor.IsotopeMz.tolist()
pqp_products = product.IsotopeMz.tolist()

for details_i, (label, ax, plotter, rt_zoom, zorder, c) in enumerate(details):
    xic_2d, xic_2d_hline, xic_2d_hline_label, ms1_chrom, chroms, rt = plotter.load_raw_data(
        pqp_precursors,
        pqp_products,
        log_heatmap=True,
        retention_time_zoom=rt_zoom,
        number_of_isotopes=n_isotopes,
        normalise_per_fragment=normalise_per_fragment,
    )
    ax.imshow(
        xic_2d,
        interpolation='none',
        extent=[min(rt), max(rt), 0, len(xic_2d)],
        aspect='auto',
        cmap=plt.cm.viridis,
    )
    
    plt.setp(ax.get_xticklabels(), visible=False)
    ax.grid(False)

    for i, hline in enumerate(reversed(xic_2d_hline[:-1])):
        is_bounding_line = i % (n_isotopes + 1) == n_isotopes
        alpha = 0.4 if is_bounding_line else 0.1
        ls = '-' if is_bounding_line else '--'
        ax.axhline(hline - 0.5, ls=ls, c='w', alpha=alpha)
    
    mz_band_count = len(pqp_precursors) + len(pqp_products)
    mz_band_height = len(xic_2d) / mz_band_count
    
    if rt_zoom is not None:
        ax.yaxis.set_ticklabels([])
    else:
        tick_labels = [f'MS2({i})' for i in reversed(range(1, mz_band_count))] + ['MS1']
        ax.set_yticks(np.arange(mz_band_height / 2, mz_band_height * mz_band_count, mz_band_height))
        ax.set_yticklabels(tick_labels, fontsize=10)
        ax.set_ylabel(label)

    # plot MS1
    ax_ms1 = chrom if rt_zoom is None else chrom_zoom
    ls = '--' if label == 'Combined' else '-'
    ax_ms1.plot(rt, ms1_chrom, ls=ls, c=c, label=label, zorder=zorder, lw=3)
    ax_ms1.set_xlim((rt[0], rt[-1]))
    ax_ms1.yaxis.set_ticklabels([])
    if details_i == 5:
        ax_ms1.legend()
        ax_ms1.set_ylabel('MS1 signal')
    ax_ms1.set_xlabel('Retention time (s)')

fig.savefig(f'{fig_dir}/fragments-{pqm}.pdf')
fig.savefig(f'{fig_dir}/fragments-{pqm}.png')
plt.show()
```
