## Supplementary material for "Toffee – a highly efficient, lossless file format for DIA-MS": Jupyter notebooks: in-silico-dilution.ipynb

```
# %% [markdown]
# In-silico Dilution of Toffee Data

With the advantages of `toffee` we can create an in-silico dilution data set for further analysis of pipelines, such as global FDR control, where we have a ground truth.

# %% [code]
%load_ext autoreload
%autoreload 2

# %% [code]
import os
import uuid

from IPython.display import SVG, display_markdown

import matplotlib.pyplot as plt

import numpy as np

import seaborn as sns

import pandas as pd

from pipeline import DevPipeline, PyProphetColumns

import toffee
from toffee.manipulation import InSilicoDilutionConstructor
from toffee.util import calculate_isotope_mz
from toffee.viz import ToffeeFragmentsPlotter

from tqdm import tqdm

import xarray as xr

base_dir = os.environ.get('DIA_TEST_DATA_REPO', None)
assert base_dir is not None
fig_ext = 'svg'

sns.set(
    palette='husl',
    font_scale=2,
    font='serif',
)

base_output_dir = 'results/in-silico-dilution/'
results_path = f'{base_output_dir}/raw'
fig_dir = 'results/figures'
os.makedirs(base_output_dir, exist_ok=True)
os.makedirs(results_path, exist_ok=True)
os.makedirs(fig_dir, exist_ok=True)

# %% [code]
toffee.__version__

# %% [markdown]
## Building the in-silico data

In general terms, we take the approach of starting with a `background` toffee file, and adding specific peptides from a `foreground` toffee file. The peptides in the foreground file have a known retention time, and we can control the *dilution* in which they are added to the background.

# %% [markdown]
### Selecting the peptides

In order to add our foreground peptides, we need the mass over charge ratio for each precursor and product ion, in addition to their respective retention time that has been found from previous analysis of the data to be used in the foreground.

We are using the 2,000 most confidently identified peptides with at least 5 fragments from a previous `ProCan90-M06-07` run. The retention times of each precursor have been previously calculated and stored on disk, this is loaded in and stored as `ForegroundRetentionTime`.

# %% [code]
def spectral_library_propterties(srl_path, psm_list_path):
    srl = pd.read_csv(srl_path, sep='\t')
    srl['PrecursorId'] = srl.ModifiedPeptideSequence + '_' + srl.PrecursorCharge.map(str)
    srl = srl[srl.Decoy == 0]
    
    psm_list = pd.read_csv(psm_list_path, sep='\t')
    srl = pd.merge(srl, psm_list, on='TransitionGroupId') \
        .rename(columns={'InSilicoRT': 'ForegroundRetentionTime'})

    print('Number of PQP:', srl.TransitionGroupId.unique().size)
    print('Number of modified peptides:', srl.ModifiedPeptideSequence.unique().size)
    print('Number of peptides:', srl.PeptideSequence.unique().size)
    print('Number of proteins:', srl.ProteinId.unique().size)

    fig, (ax1, ax2) = plt.subplots(ncols=2, figsize=(15, 6))
    
    ax = ax1
    df = srl.drop_duplicates(subset=['ProteinId', 'PeptideSequence'])
    df = df.PeptideSequence.groupby(df.ProteinId).count()
    df.hist(bins=df.max(), ax=ax)
    ax.set_xlabel('No. of Peptides per Protein')
    ax.set_ylabel('Frequency')
    
    ax = ax2
    psm_list.InSilicoRT.hist(bins=np.linspace(0, 90 * 60, 11), ax=ax)
    ax.set_xlabel('Expected Retention Time (s)')
    ax.set_ylabel('Frequency')
    
    fig.tight_layout()
    plt.show()
    return srl

# %% [code]
spectral_library_fnames = {
    'Simple': base_dir + '/in-silico-dilution/srl/hek-top2000.with_decoys.tsv',
    'Complex': base_dir + '/in-silico-dilution/srl/hek-top2000.ecoli.with_decoys.tsv',
}
spectral_library_psm_list_fname = base_dir + '/in-silico-dilution/srl/hek-top2000.retention_times.tsv.gz'
spectral_library = {}
for k, fname in spectral_library_fnames.items():
    print('-' * 10, k)
    spectral_library[k] = spectral_library_propterties(fname, spectral_library_psm_list_fname)

# %% [code]
irt_library_fnames = {
    'Simple': base_dir + '/in-silico-dilution/srl/hek-top2000.iRT.tsv',
    'Complex': base_dir + '/in-silico-dilution/srl/hek-top2000.ecoli.iRT.tsv',
}
irt_library_psm_list_fname = base_dir + '/in-silico-dilution/srl/hek-top2000.irt.retention_times.tsv.gz'
irt_library = {}
for k, fname in irt_library_fnames.items():
    print('-' * 10, k)
    irt_library[k] = spectral_library_propterties(fname, irt_library_psm_list_fname)

# %% [code]
expected_retention_time_df = spectral_library['Simple'].rename(columns={'TransitionGroupId': 'PQM'})
expected_retention_time_df = expected_retention_time_df[['PQM', 'ForegroundRetentionTime']]
expected_retention_time_df = expected_retention_time_df.drop_duplicates()
expected_retention_time_df.set_index('PQM', inplace=True)

# %% [markdown]
### Build the dilutions

We are going to create two dilution series, one with a blank background and another with a yeast background. The former will show us the best case scenario where the software should be able to detect all precursors that are above the limit of detection, while the second will show us the impact of noise on the software's ability to accurately detect precursors in amongst noise.

It is worth noting that these dilution series cannot model the true noise of a mass spectrometer. In particular, the additive noise cannot be captured with this method.

# %% [code]
def create_metadata():
    backgrounds = {
        'Blank': '180508_blank_s_m02_1',
        'Ecoli': '190109_Missing-1-Bcontrol-RT_007V1_007WJ_M05_S_1',
    }
    log2_dilutions = range(6)
    srls = ['Simple', 'Complex']
    data = []
    for srl in srls:
        for b, f in backgrounds.items():
            for log2_dilution in log2_dilutions:
                data.append({
                    'Dilution': 2 ** log2_dilution,
                    'Background': b,
                    'BackgroundFile': f,
                    'SRL': srl,
                })
    data = pd.DataFrame(data)

    def injection_name(row):
        return 'In-Silico-Dilution.ProCan90-M06-07.{}.{:02d}'.format(
            row.Background,
            row.Dilution,
        )

    data['InjectionName'] = data.apply(injection_name, axis=1)
    data['InjectionName'] = data.InjectionName.str.lower()
    return data

# %% [code]
row_metadata = create_metadata()

raw_data_dir = 'raw_data/in-silico-dilution/'
os.makedirs(raw_data_dir, exist_ok=True)

metadata = row_metadata.copy()
metadata['InjectionFilePath'] = f'{raw_data_dir}/' + metadata.InjectionName + '.tof'
metadata['ObservationId'] = metadata.InjectionName + '.' + metadata.SRL.str.lower()
metadata['BackgroundFilePath'] = f'{base_dir}/in-silico-dilution/tof/' + metadata.BackgroundFile + '.tof'
metadata['ForegroundFilePath'] = f'{base_dir}/in-silico-dilution/tof/ProCan90-M06-07.tof'
metadata['AddIrtFragments'] = metadata.Background != 'Ecoli'

row_metadata = row_metadata.drop('SRL', axis=1).drop_duplicates().set_index('InjectionName')
metadata.set_index('InjectionName', inplace=True)

# %% [code]
constructors = {}
for _, row in metadata.iterrows():
    if row.Background not in constructors:
        print(f'Creating InSilicoDilutionConstructor for {row.Background}', flush=True)
        if row.AddIrtFragments:
            irt_precursor_and_product_df = irt_library[row.SRL]
        else:
            irt_precursor_and_product_df = None
        constructors[row.Background] = InSilicoDilutionConstructor(
            background_tof_fname=row.BackgroundFilePath,
            foreground_tof_fname=row.ForegroundFilePath,
            precursor_and_product_df=spectral_library[row.SRL],
            irt_precursor_and_product_df=irt_precursor_and_product_df,
            show_progress_bar=False,
        )

for _, row in tqdm(metadata.iterrows()):
    if not os.path.isfile(row.InjectionFilePath):
        print(f'Creating {row.name}', flush=True)
        constructor = constructors[row.Background]
        constructor.run(
            row.InjectionFilePath,
            dilution=row.Dilution,
        )

# %% [markdown]
Let's have a look at how the dilution data is constructed. The following plots show the foreground data, original blank background, and the result for both blank and yeast backgrounds.

# %% [code]
lower_mz_overlap = 0.0
upper_mz_overlap = 1.0

old_plotter = ToffeeFragmentsPlotter(
    constructors['Ecoli'].foreground_tof_fname,
    lower_mz_overlap=lower_mz_overlap,
    upper_mz_overlap=upper_mz_overlap,
)
old_ecoli_plotter = ToffeeFragmentsPlotter(
    constructors['Ecoli'].background_tof_fname,
    lower_mz_overlap=lower_mz_overlap,
    upper_mz_overlap=upper_mz_overlap,
)
old_blank_plotter = ToffeeFragmentsPlotter(
    constructors['Blank'].background_tof_fname,
    lower_mz_overlap=lower_mz_overlap,
    upper_mz_overlap=upper_mz_overlap,
)

# %% [code]
plotter_dilution = 1
tof_fname_fmt = f'{raw_data_dir}/in-silico-dilution.procan90-m06-07.{{}}.{{:02d}}.tof'
blank_plotter = ToffeeFragmentsPlotter(
    tof_fname_fmt.format('blank', plotter_dilution),
    lower_mz_overlap=lower_mz_overlap,
    upper_mz_overlap=upper_mz_overlap,
)
ecoli_plotter = ToffeeFragmentsPlotter(
    tof_fname_fmt.format('ecoli', plotter_dilution),
    lower_mz_overlap=lower_mz_overlap,
    upper_mz_overlap=upper_mz_overlap,
)

# %% [code]
# pqm = 'AAFTAFEEAQLPR_2'
pqm = 'SKPGAAMVEMADGYAVDR_3'  # top_results.peptide_group_label.iloc[1]

n_isotopes = 0
precursor, product = calculate_isotope_mz(
    precursor_and_product_df=spectral_library['Simple'],
    n_isotopes=n_isotopes,
    transition_group_id=pqm,
    sort_by_intensity=True,
)

# %% [code]
normalise_per_fragment = False
import plotly.io as pio

output_fname = f'{fig_dir}/fragments.original.{pqm}.{fig_ext}'
fig = old_plotter.create_plotly_figure(
    precursor.IsotopeMz.tolist(),
    product.IsotopeMz.tolist(),
    'Foreground HEK293: ' + pqm,
    number_of_isotopes=n_isotopes,
    normalise_per_fragment=normalise_per_fragment,
    output_fname=output_fname,
)
if fig_ext == 'svg':
    display(SVG(filename=output_fname))
    pio.write_image(fig, output_fname.replace(fig_ext, 'pdf'))

output_fname = f'{fig_dir}/fragments.original_blank.{pqm}.{fig_ext}'
fig = old_blank_plotter.create_plotly_figure(
    precursor.IsotopeMz.tolist(),
    product.IsotopeMz.tolist(),
    'Background Blank: ' + pqm,
    number_of_isotopes=n_isotopes,
    normalise_per_fragment=normalise_per_fragment,
    output_fname=output_fname,
)
if fig_ext == 'svg':
    display(SVG(filename=output_fname))
    pio.write_image(fig, output_fname.replace(fig_ext, 'pdf'))

output_fname = f'{fig_dir}/fragments.blank_{plotter_dilution:04d}.{pqm}.{fig_ext}'
fig = blank_plotter.create_plotly_figure(
    precursor.IsotopeMz.tolist(),
    product.IsotopeMz.tolist(),
    'Blank: ' + pqm,
    number_of_isotopes=n_isotopes,
    normalise_per_fragment=normalise_per_fragment,
    output_fname=output_fname,
)
if fig_ext == 'svg':
    display(SVG(filename=output_fname))
    pio.write_image(fig, output_fname.replace(fig_ext, 'pdf'))

output_fname = f'{fig_dir}/fragments.original_ecoli.{pqm}.{fig_ext}'
fig = old_ecoli_plotter.create_plotly_figure(
    precursor.IsotopeMz.tolist(),
    product.IsotopeMz.tolist(),
    'Background E.coli: ' + pqm,
    number_of_isotopes=n_isotopes,
    normalise_per_fragment=normalise_per_fragment,
    output_fname=output_fname,
)
if fig_ext == 'svg':
    display(SVG(filename=output_fname))
    pio.write_image(fig, output_fname.replace(fig_ext, 'pdf'))

output_fname = f'{fig_dir}/fragments.ecoli_{plotter_dilution:04d}.{pqm}.{fig_ext}'
fig = ecoli_plotter.create_plotly_figure(
    precursor.IsotopeMz.tolist(),
    product.IsotopeMz.tolist(),
    'E.coli + HEK293: ' + pqm,
    number_of_isotopes=n_isotopes,
    normalise_per_fragment=normalise_per_fragment,
    output_fname=output_fname,
)
if fig_ext == 'svg':
    display(SVG(filename=output_fname))
    pio.write_image(fig, output_fname.replace(fig_ext, 'pdf'))

# %% [markdown]
## Run the pipeline

# %% [code]
swath_scores_kwargs = {
    'Simple': {
        'rt_transform_path': f'{base_dir}/in-silico-dilution/srl/hek-top2000.trafoXML',
        'specimen_suffix': '.simple',
    },
    'Complex': {
        'irt_srl_path': irt_library_fnames['Complex'],
        'specimen_suffix': '.complex',
    },
}
result_data_keys = [
    PyProphetColumns.INTENSITY,
    PyProphetColumns.RETENTION_TIME,
    PyProphetColumns.M_SCORE,
]

unmerged_batch_xarray = {}
for srl in metadata.SRL.unique():
    print('-' * 10, srl, flush=True)
    batch_pipeline = DevPipeline.default_open_ms_toffee_batch_pipeline(
        srl_path=spectral_library_fnames[srl].replace('.tsv', '.pqp'),
        swath_scores_kwargs=swath_scores_kwargs[srl],
    )
    unmerged_batch_xarray[srl] = batch_pipeline.run(
        metadata_df=metadata.loc[metadata.SRL == srl],
        results_path=results_path,
        swath_scores_kwargs={
            'minimal_quality': -5,
        },
        fdr_kwargs={
            'run_name': srl,
            'apply_protein_fdr': False,  # doesn't make sense for in-silico data
            'pi0_lambda': (0.0, 0.0001, 0.00001),
        },
        result_data_keys=result_data_keys,
    )

# %% [code]
def stack_xarray(unmerged_xarray, row_metadata):
    dataset = []
    for k, v in unmerged_xarray.items():
        v = v.rename({'InjectionName': 'InjectionNameId'}).expand_dims('SpectralLibrary')
        v['SpectralLibrary'] = ('SpectralLibrary', [k])
        
        # split the FDR iterations
        ids = v['InjectionNameId'].values
        v.coords['InjectionName'] = (
            'InjectionNameId',
            list(map(lambda x: '-'.join(x.split('-')[:-1]), ids)),
        )
        v.coords['FDRIter'] = (
            'InjectionNameId',
            list(map(lambda x: int(x.split('-')[-1]), ids)),
        )
        
        # add the metadata
        for c in row_metadata.columns:
            v.coords[c] = (
                'InjectionNameId',
                row_metadata.loc[v['InjectionName'], c],
            )
            
        dataset.append(v)
        
    dataset = xr.concat(dataset, dim='SpectralLibrary')
            
    # rename the RT DataArray
    dataset = dataset.rename({'RT': 'RetentionTime'})

    # set the confidence DataArray
    mask = dataset['m_score'].values == 0
    dataset['m_score'].values[mask] = 1e-15
    dataset['Confidence'] = (-np.log10(dataset['m_score']) - 2) / 13
    dataset = dataset.drop('m_score')

    return dataset

ds = stack_xarray(unmerged_batch_xarray, row_metadata)
ds.to_netcdf(f'{results_path}/in-silico-dilution.nc')

# %% [markdown]
## Manipulating the output

Using the raw data returned, we can also normalise the intensities:

# %% [code]
ds = xr.open_dataset(f'{results_path}/in-silico-dilution.nc')
ds

# %% [code]
intensity = ds['Intensity'].to_dataframe().reset_index()
intensity[~intensity.Intensity.isna()].head()

# %% [code]
def normalise_intensities(xarr, global_filter_funcs, normalisation_filter_funcs):
    """
    Normalise the intensities based on data within in the `xarr` argument.
    
    :param xarr (xarray.Dataset): a properly formed Dove xarray object
    :param global_filter_funcs (list[functor]): a list of functors that expect an xarray
        DataArray and return a boolean mask that is the same shape as the input array. These
        functors to filter the input array down to the data that should be normalised
    :param normalisation_filter_funcs (list[functor]): a list of functors that expect an xarray
        DataArray and return a boolean mask that is the same shape as the input array. These
        functors to filter the output of the `global_filter_funcs` filtering to the vector(s)
        that will be used for calculating the normalising vector
    
    :return (xarray.Dataset): a new Dove xarray that appends a new page to the input called
        'NormalisedIntensity'. For example,
    
        >>> global_filter_funcs = [
        >>>     lambda x: x.Background == 'Blank',
        >>> ]
        >>> normalisation_filter_funcs = [
        >>>     lambda x: x.SpectralLibrary == 'Simple',
        >>>     lambda x: x.Dilution == 1,
        >>> ]
        >>> new_xarr = normalise_intensities(xarr, global_filter_funcs, normalisation_filter_funcs)
        >>> assert np.all(new_xarr.Background == 'Blank')
        >>> assert np.any(new_xarr.SpectralLibrary != 'Simple')
        >>> assert np.any(new_xarr.Dilution != 1)
    """
    xarr = xarr.copy()
    
    def filter_xarray(xarr, filter_funcs):
        mask = None
        for func in filter_funcs:
            tmp_mask = func(xarr)
            mask = tmp_mask if mask is None else mask & tmp_mask
        return xarr if mask is None else xarr.where(mask, drop=True)

    filtered_ij = filter_xarray(xarr.Intensity, global_filter_funcs)
    norm_a_ij = filter_xarray(filtered_ij, normalisation_filter_funcs)

    pqm_nan_mask = norm_a_ij.notnull().all(dim='InjectionNameId').all(dim='SpectralLibrary')
    norm_b_ij = norm_a_ij.sel(PQM=pqm_nan_mask)
    assert norm_b_ij.notnull().all()

    tic_i = norm_b_ij.sum(dim='PQM')
    tic_ave = tic_i.mean(dim='InjectionNameId')
    scale_i = tic_ave / tic_i

    norm_c_ij = scale_i * norm_a_ij

    norm_d_j = norm_c_ij.median(dim='InjectionNameId')
    if norm_d_j.SpectralLibrary.shape[0] != xarr.SpectralLibrary.shape[0]:
        norm_d_j_list = []
        for srl in xarr.SpectralLibrary.values:
            norm_d_j_list.append(norm_d_j.assign_coords(SpectralLibrary=[srl]))
        norm_d_j = xr.concat(norm_d_j_list, dim='SpectralLibrary')

    xarr['NormalisedIntensity'] = filtered_ij / norm_d_j
    return xarr

# %% [code]
global_filter_funcs = [
]
normalisation_filter_funcs = [
    lambda x: x.SpectralLibrary == 'Simple',
    lambda x: x.Background == 'Blank',
    lambda x: x.Dilution == 1,
]

ds = normalise_intensities(ds, global_filter_funcs, normalisation_filter_funcs)

# %% [markdown]
and calculate the offsets from expected retention time:

# %% [code]
def calculate_delta_rt(xarr, expected_retention_time_df):
    """
    :param xarr (xarray.Dataset): a properly formed Dove xarray object
    :param expected_retention_time_df (pd.DataFrame): a dataframe with PSMs as the index, and
        a column named 'ExpectedRT'.
    
    :return (xarray.Dataset): a new Dove xarray that appends a new page to the input called
        'AbsDeltaRetentionTime'
    """
    xarr = xarr.copy()
    expected_rt = xr.DataArray(
        expected_retention_time_df.reindex(xarr.PQM.values).ForegroundRetentionTime,
    )
    assert xarr.PQM.shape[0] == expected_rt.PQM.shape[0]
    xarr['AbsDeltaRetentionTime'] = np.abs(xarr['RetentionTime'] - expected_rt)
    return xarr
ds = calculate_delta_rt(ds, expected_retention_time_df)

# %% [code]
ds.to_netcdf(f'{results_path}/in-silico-dilution.full.nc')

# %% [markdown]
## Results
Now, we can plot the results

# %% [code]
ds = xr.open_dataset(f'{results_path}/in-silico-dilution.full.nc')
ds

# %% [markdown]
By normalising against the original intensity, we can assess how closely the peaks match to their theoretical dilution

- Blank are lower because of the limit of detection of the instrument (i.e. no individual detection can be lower than 36)
- Yeast does better with dilution, however, this is at the expense of over-estimating intensity in some situations suggesting that we are quantifying noise in some way

# %% [code]
palette = [
    plt.cm.Blues_r(0.2),
    plt.cm.Reds_r(0.1),
]

# %% [code]
def plot_intensity(xarr, expected_retention_time_df):
    col = 'NormalisedIntensity'
    df = xarr[col].to_dataframe().reset_index(drop=False)
    df = df.set_index('PQM').loc[expected_retention_time_df.index].reset_index(drop=False)
    df = df.loc[df.FDRIter == 0]
    
    new_col = 'Rel Log2 Intensity'
    df[new_col] = df[col].apply(np.log2)

    col_order = ['Simple', 'Complex']
    hue_order = ['Blank', 'Ecoli']
#     palette = 'viridis'#'husl'
    for use_box in [True, False]:
        fig = plt.Figure()
        if use_box:
            kwargs = dict(
                kind='box',
            )
        else:
            kwargs = dict(
                kind='strip',
                jitter=0.25,
                dodge=True,
                edgecolor='white',
            )
        fg = sns.catplot(
            data=df,
            x='Dilution',
            y=new_col,
            col='SpectralLibrary',
            hue='Background',
            col_order=col_order,
            hue_order=hue_order,
            palette=palette,
            **kwargs,
        )
        dilutions = df.Dilution.unique()
        for ax in fg.axes.flatten():
            for dilution, loc in zip(dilutions, ax.get_xaxis().get_majorticklocs()):
                y = np.log2(1.0 / dilution)
                ax.plot([loc - 0.5, loc + 0.5], [y, y], 'r', zorder=100)
        plt.show()

# %% [code]
plot_intensity(ds, expected_retention_time_df)

# %% [markdown]
What about retention time deltas

# %% [code]
def plot_rt(xarr, expected_retention_time_df, false_positive_threshold=10, delta_rt_lower_bound=1e-2):
    col = 'AbsDeltaRetentionTime'
    df = xarr[col].to_dataframe().reset_index(drop=False)
    df = df.set_index('PQM').loc[expected_retention_time_df.index].reset_index(drop=False)
    df = df.loc[df.FDRIter == 0]

    df.loc[df[col] <= delta_rt_lower_bound, col] = delta_rt_lower_bound
    
    new_col = col + 'Log10'
    df[new_col] = df[col].apply(np.log10)

    col_order = ['Simple', 'Complex']
    hue_order = ['Blank', 'Ecoli']
#     palette = 'viridis'#'husl'
    fg = sns.catplot(
        kind='strip',
        data=df,
        x='Dilution',
        y=new_col,
        col='SpectralLibrary',
        hue='Background',
        col_order=col_order,
        hue_order=hue_order,
        palette=palette,
        edgecolor='white',
        jitter=True,
        dodge=True,
    )
    for ax in fg.axes.flatten():
        for loc in ax.get_xaxis().get_majorticklocs():
            y = np.log10(false_positive_threshold)
            ax.plot([loc - 0.5, loc + 0.5], [y, y], 'r', zorder=100)

# %% [code]
plot_rt(ds, expected_retention_time_df)

# %% [markdown]
## Combined figure

# %% [code]
import matplotlib.gridspec as gridspec

fig, ax = plt.subplots(figsize=(20, 13))

gs = gridspec.GridSpec(3, 4)
gs.update(wspace=0.55, hspace=0.45)

ax_intensity = plt.subplot(gs[0, :])

ax_tp = plt.subplot(gs[1, :2])
ax_fp = plt.subplot(gs[2, :2], sharex=ax_tp)
ax_fn = plt.subplot(gs[2, 2:], sharex=ax_tp)

palette = [
    plt.cm.Blues_r(0.1),
    plt.cm.Reds_r(0.1),
    plt.cm.Blues_r(0.4),
    plt.cm.Reds_r(0.4),
]
hue_order = [
    'Simple SRL / Water Background',
    'Complex SRL / Water Background',
    'Simple SRL / Ecoli Background',
    'Complex SRL / Ecoli Background',
]

kwargs = dict(
    xy=(-50, 200),
    xycoords='axes points',
    va='center',
    ha='left',
    fontsize=20,
    weight='bold',
)
label_fontsize = 16

ax_intensity.annotate('A.', **kwargs)
ax_tp.annotate('B.', **kwargs)
ax_fp.annotate('C.', **kwargs)
ax_fn.annotate('D.', **kwargs)

# ******************
col = 'NormalisedIntensity'
df = ds[col].to_dataframe().reset_index(drop=False)
df = df.set_index('PQM').loc[expected_retention_time_df.index].reset_index(drop=False)
df = df.loc[df.FDRIter == 0]

new_col = 'Normalised Intensity (Log2)'
df[new_col] = df[col].apply(np.log2)

df.loc[df.Background == 'Blank', 'Background'] = 'Water'
df['Label'] = df.SpectralLibrary + ' SRL / ' + df.Background + ' Background'

ax = ax_intensity
sns.stripplot(
    data=df,
    x='Dilution',
    y=new_col,
    hue='Label',
    hue_order=hue_order,
    palette=palette,
    jitter=0.25,
    dodge=True,
    edgecolor='white',
    ax=ax,
)
dilutions = df.Dilution.unique()
for dilution, loc in zip(dilutions, ax.get_xaxis().get_majorticklocs()):
    y = np.log2(1.0 / dilution)
    ax.plot([loc - 0.5, loc + 0.5], [y, y], 'r', zorder=100)
ax.legend_ = None
ax.set_title('Normalised Intensity')
ax.set_ylabel('Log2', fontsize=label_fontsize)
ax.set_xlabel('Dilution', fontsize=label_fontsize)

# ******************
filter_size_number_of_seconds = 10
col = 'AbsDeltaRetentionTime'
df = ds[col].to_dataframe()[[col, 'Background', 'Dilution']].reset_index(drop=False)
df['InjectionName'] = df.InjectionNameId.apply(lambda x: '-'.join(x.split('-')[:-1]))
df['FDRIter'] = df.InjectionNameId.apply(lambda x: int(x.split('-')[-1]))
df = df.set_index(['SpectralLibrary', 'InjectionName', 'FDRIter', 'Background', 'Dilution', 'PQM'])
df = df.drop('InjectionNameId', axis=1)
df = df.unstack()
df = df.dropna(how='all', axis=1)
num_psms = df.shape[1]
df = pd.DataFrame(
    {
        'Number of PSMs': num_psms - df.isna().sum(axis=1),
        'False Positives': (df > filter_size_number_of_seconds).sum(axis=1),
    },
    index=df.index,
)
df.reset_index(drop=False, inplace=True)

df['Percentage of PSMs'] = 100 * df['Number of PSMs'] / num_psms
df['True Positives'] = df['Number of PSMs'] - df['False Positives']
df['False Discovery Rate'] = 100 * df['False Positives'] / df['Number of PSMs']

# compare false negatives to the number of true positives found in the easiest
# run: blank + simple SRL
df['False Negatives'] = pd.np.nan
for dilution in df.Dilution.unique():
    num_expected_psms = df.loc[
        (df.Dilution == dilution) & 
        (df.Background == 'Blank') & 
        (df.SpectralLibrary == 'Simple'),
        'True Positives'
    ].values[0]
    mask = df.Dilution == dilution
    df.loc[mask, 'False Negatives'] = num_expected_psms - df.loc[mask, 'True Positives']
df['False Negative Rate'] = 100 * df['False Negatives'] / (df['False Negatives'] + df['True Positives'])

df.loc[df.Background == 'Blank', 'Background'] = 'Water'
df['Label'] = df.SpectralLibrary + ' SRL / ' + df.Background + ' Background'

ax = ax_tp
sns.barplot(
    data=df,
    x='Dilution',
    y='True Positives',
    hue='Label',
    hue_order=hue_order,
    palette=palette,
    ax=ax,
)
ax.legend(bbox_to_anchor=(1.15, 0.5), loc=6, borderaxespad=0.0, frameon=False)
ax.set_title(ax.yaxis.get_label().get_text())
ax.set_ylabel('Count', fontsize=label_fontsize)
ax.set_xlabel('Dilution', fontsize=label_fontsize)

ax = ax_fp
sns.barplot(
    data=df,
    x='Dilution',
    y='False Positives',
    hue='Label',
    hue_order=hue_order,
    palette=palette,
    ax=ax,
)
ax.legend_ = None
ax.set_title(ax.yaxis.get_label().get_text())
ax.set_ylabel('Count', fontsize=label_fontsize)
ax.set_xlabel('Dilution', fontsize=label_fontsize)

ax = ax_fn
sns.barplot(
    data=df,
    x='Dilution',
    y='False Negatives',
    hue='Label',
    hue_order=hue_order,
    palette=palette,
    ax=ax,
)
ax.legend_ = None
ax.set_title(ax.yaxis.get_label().get_text())
ax.set_ylabel('Count', fontsize=label_fontsize)
ax.set_xlabel('Dilution', fontsize=label_fontsize)

# ******************
fig.savefig(f'{fig_dir}/isd-confusion-matrix.pdf')
plt.show()

# %% [code]
```
