## Supplementary material for "Toffee – a highly efficient, lossless file format for DIA-MS": Jupyter notebooks: missing-fragment-visualisation.ipynb

```
# %% [markdown]
# Investigating the Missing PQMs

Through the `ToffeeFragmentsPlotter` we can start to explore the raw data contained within a toffee file. These plots are specifically aimed at extracting the same data that one might use when calculating scores in a tool such as `OpenMSToffee`.

# %% [code]
%load_ext autoreload
%autoreload 2

# %% [code]
import os

from IPython.display import SVG, display_markdown

import pandas as pd

import toffee
from toffee.util import calculate_isotope_mz
from toffee.viz import ToffeeFragmentsPlotter

# %% [code]
toffee.__version__

# %% [code]
def plot_fragments(experiment, specimen, nplots=2):
    # use a spectral library to define the precursor/product mass over charge ratios
    if experiment == 'roest2016':
        srl_fname = 'srl/roest2016/SpyogenesAssayLibrary.with_decoys.tsv'
    elif experiment == 'procan90':
        srl_fname = 'srl/procan90/hek_srl.OpenSwath.reverse_decoys.tsv'
    else:
        raise VallueError(f'Invalid experiment: {experiment}')
        
    srl = pd.read_csv(srl_fname, sep='\t').drop_duplicates()
    srl['TransitionGroupId_bkp'] = srl.TransitionGroupId
    srl['TransitionGroupId'] = srl.ModifiedPeptideSequence + '_' + srl.PrecursorCharge.map(str)
    
    #  calculate the C12/C13 isotope offsets and use these in our visualisation
    n_isotopes = 1
    precursor, product = calculate_isotope_mz(
        srl,
        n_isotopes=n_isotopes,
        sort_by_intensity=True,
    )
    precursor.set_index('Id', inplace=True)
    product.set_index('GroupId', inplace=True)
    
    # construct the plotter
    plotter = ToffeeFragmentsPlotter(
        f'raw_data/{experiment}/{specimen}.e.tof',
        use_ms1=True,
        extraction_width=20,
        lower_mz_overlap=0.0,
        upper_mz_overlap=1.0,
    )
    
    # create plots for each of the missing types
    file_types = [
        'Centroid',
        'Toffee',
        'mzML',
        'Profile',
    ]
    for file_type in file_types:
        display_markdown(f'### Missing {file_type} PQMs', raw=True)

        missing_pqms_fname = f'missing-{file_type.lower()}-pqms.tsv'
        df = pd.read_csv(f'results/{experiment}/{missing_pqms_fname}', sep='\t')
        
        ctr = 0
        for _, row in df.iterrows():
            if row.InjectionName != specimen:
                continue
            transition_group = row.PQM
            precursors = precursor.loc[transition_group].IsotopeMz.tolist()
            products = product.loc[transition_group].IsotopeMz.tolist()[:6 * (1 + n_isotopes)]

            rt_width = 300
            title = f'{transition_group} || Missing in: {row.NumberOfMissingFiles}, with q-value = {row.MedianQValue:.2e}'
            output_fname = f'results/{experiment}/{specimen}_{transition_group}.svg'
            figure = plotter.create_plotly_figure(
                precursors,
                products,
                psm_name=title,
                number_of_isotopes=n_isotopes,
                retention_time_zoom=(row.RetentionTimeMedian - rt_width, row.RetentionTimeMedian + rt_width),
                retention_time_line=row.RetentionTimeMedian,
                aspect_scale=0.1,
                output_fname=output_fname,
            )
            display(SVG(filename=output_fname))

            ctr += 1
            if ctr == nplots:
                break

# %% [markdown]
## ProCan90 data

# %% [code]
plot_fragments(
    experiment='procan90',
    specimen='ProCan90-M01-01',
)

# %% [markdown]
## Roest2016 data

# %% [code]
plot_fragments(
    experiment='roest2016',
    specimen='Strep10Repl1_R03_SW',
)

# %% [code]
```
