## Supplementary material for "Toffee – a highly efficient, lossless file format for DIA-MS": Jupyter notebooks: random-access-timing.ipynb

```
# %% [markdown]
# Timing raw data access

The following analysis code shows a comparison of scan-by-scan spectrum access, and peptide-centric XIC access of `mzML` and `toffee` files. Spectrum access is the best case scenario for the `mzML` file as the file index enables direct location of the relevant spectra in the file. Peptide-centric access refers to taking a slice of the raw data across all retention time for a constrained m/z range.

It is worth noting, that these types of timings are somewhat artificial. The best method of loading and caching raw data will be highly dependent on the application, algorithm, and its implementation -- performance is always best achieved through profiling the actual code being optimized. Furthermore, there is no consideration given here to threaded access, or high memory environments that would allow all data to be held in RAM.

# %% [code]
%matplotlib inline
%load_ext autoreload
%autoreload 2

import datetime
import gc
import json
import logging
import os
from collections import namedtuple

import matplotlib.pyplot as plt

import numpy as np

import pandas as pd

import pyteomics
from pyteomics import mzml

import seaborn as sns

import toffee
from toffee.log import set_stream_logger

sns.set(
    font_scale=1,
    font='serif',
    palette='husl',
)

# %% [code]
print(' Analysis date:', datetime.datetime.now())
print('toffee version:', toffee.__version__)

# %% [code]
# filter SRL to only include window ms2-050
ms2_name = 'ms2-050'
lower_filter, upper_filter = 653.5, 659.5

base_dir = os.environ.get('DIA_TEST_DATA_REPO', None)
srl_fname = f'{base_dir}/in-silico-dilution/srl/hek-top2000.with_decoys.tsv'
srl = pd.read_csv(srl_fname, sep='\t', engine='c', low_memory=False)
srl = srl.loc[~srl.PeptideGroupLabel.str.startswith('DECOY')]

mask1 = srl.PrecursorMz >= lower_filter
mask2 = srl.PrecursorMz < upper_filter
srl = srl.loc[mask1 & mask2]

mz_to_extract = []
for precursor_mz, g in srl.groupby('PrecursorMz'):
    mz_to_extract.append((precursor_mz, g.ProductMz.tolist()))

# %% [markdown]
## Helper classes

In order to make these comparisons, and use the jupyter timer, we need to create a couple of helper classes. The `mzML` reader is slightly more complex, in that we need to write some logic that enables peptide-centric access to the spectra. 

# %% [code]
class MzMLReader():
    Window = namedtuple(
        'Window',
        ['name', 'center', 'lower', 'upper', 'ids'],
    )

    def __init__(
        self,
        mzml_fname,
        debug=False,
    ):
        mzml_ext = os.path.splitext(mzml_fname)[1].lower()
        if mzml_ext != '.mzml':
            raise ValueError(f'Invalid input file extension (should be .mzml): {mzml_ext}')
        if not os.path.isfile(mzml_fname):
            raise ValueError(f'Input file does not exist: {mzml_fname}')
        self.mzml_fname = mzml_fname

        logger_kwargs = {}
        if debug:
            logger_kwargs['level'] = logging.DEBUG
        self.logger = set_stream_logger(**logger_kwargs)
        self.debug = debug

        self._ms1_window = self.Window('ms1', -1, -1, -1, [])
        self._ms2_windows = []

    def setup_windows(self):
        self.logger.debug('Collecting the window locations in the mzML file')
        with mzml.PreIndexedMzML(self.mzml_fname) as reader:
            for i, ele in enumerate(reader):
                precursor_list = ele.get('precursorList', {})
                ms_level = 1 if not precursor_list else 2
                ms_level = ele.get('ms level', ms_level)
                if ms_level is None:
                    msg = json.dumps(ele, indent=2, cls=_NumpyEncoder)
                    raise ValueError(f'No MS level found!\n{msg}')

                scan_id = ele.get('id', None)
                if scan_id is None:
                    msg = json.dumps(ele, indent=2, cls=_NumpyEncoder)
                    raise ValueError(f'No id found!\n{msg}')

                if ms_level == 1:
                    window = self._ms1_window
                else:
                    window = self._get_ms2_window(ele)
                window.ids.append(scan_id)
        
    def extract_spectra(
        self,
        use_ms1,
        n=1000,
        random_seed=0,
    ):
        np.random.seed(random_seed)
        if len(self._ms1_window.ids) == 0 or len(self._ms2_windows) != 100:
            raise RuntimeError('Ensure you have run self.setup_windows()')
            
        if use_ms1:
            window = self._ms1_window
        else:
            window = self._ms2_windows[49]  # ms2-050
            
        n_spectra = len(window.ids)
        spectra_indices = np.random.randint(low=0, high=n_spectra - 1, size=(n,))
        output = []
        with mzml.PreIndexedMzML(self.mzml_fname) as reader:
            for idx in spectra_indices:
                scan_id = window.ids[idx]
                _, mz, intensity = self._get_spectra(reader, scan_id)
                output.append((mz, intensity))

        # force the garbage collector to manage memory more explicitly
        gc.collect()

        return output
        
    def extract_xic(
        self,
        precursor_mz,
        product_mz_list,
        use_ms1=True,
        ppm_extraction_width=50.0,
        lower_window_overlap=0.0,
        upper_window_overlap=1.0,
    ):
        if len(self._ms2_windows) == 0:
            raise RuntimeError('Ensure you have run self.setup_windows()')
            
        xic = []
        if use_ms1:
            self.logger.debug('Extract precursor XIC from MS1 window')
            xic.extend(self._extract_xic(self._ms1_window, [precursor_mz], ppm_extraction_width))
            
        self.logger.debug('Extract product XICs from the MS2 window')
        window = None
        for w in self._ms2_windows:
            assert isinstance(w, self.Window)
            if w.lower + lower_window_overlap <= precursor_mz < w.upper - upper_window_overlap:
                window = w
                break
        if window is None:
            raise RuntimeError(f'Could not find MS2 window for {precursor_mz}')
        xic.extend(self._extract_xic(window, product_mz_list, ppm_extraction_width))

        return xic

    def _get_ms2_window(self, ele):
        """
        MS2 windows are centered around a common m/z value. Use this to make sure that new id's are added
        to the same window
        """
        precursor_list = ele.get('precursorList', {})
        num_precursors = precursor_list.get('count', 0)
        if num_precursors != 1:
            msg = json.dumps(ele, indent=2, cls=_NumpyEncoder)
            raise ValueError(f'There should only be one precursor!\n{msg}')
        precursor = precursor_list.get('precursor')[0]
        isolation = precursor.get('isolationWindow', {})
        center = isolation.get('isolation window target m/z', None)
        if center is None:
            msg = json.dumps(ele, indent=2, cls=_NumpyEncoder)
            raise ValueError(f'No window center found!\n{msg}')
        window = None
        for w in self._ms2_windows:
            if toffee.MathUtils.almostEqual(w.center, center):
                window = w
                break
        if window is None:
            lower_offet = isolation.get('isolation window lower offset', None)
            upper_offet = isolation.get('isolation window upper offset', None)
            window_id = len(self._ms2_windows) + 1
            window_name = toffee.ToffeeWriter.ms2Name(window_id)
            window = self.Window(
                window_name,
                center,
                center - lower_offet,
                center + upper_offet,
                [],
            )
            self._ms2_windows.append(window)
        return window

    def _extract_xic(self, window, product_mz_list, ppm_extraction_width):
        assert isinstance(window, self.Window)
        
        num_products = len(product_mz_list)
        num_rt = len(window.ids)
        output = []
        for i in range(num_products):
            output.append((np.zeros(num_rt), np.zeros(num_rt)))

        # add raw data to the scan
        with mzml.PreIndexedMzML(self.mzml_fname) as reader:
            for scan_idx, scan_id in enumerate(window.ids):
                scan_rt, mz, intensity = self._get_spectra(reader, scan_id)
                for product_idx in range(num_products):
                    output[product_idx][0][scan_idx] = scan_rt

                    product_mz = product_mz_list[product_idx]
                    mz_offset = ppm_extraction_width * 0.5e-6 * product_mz
                    start_idx = np.searchsorted(mz, product_mz - mz_offset)
                    end_idx = np.searchsorted(mz, product_mz + mz_offset)
                    output[product_idx][1][scan_idx] = np.sum(intensity[start_idx:end_idx])

        # force the garbage collector to manage memory more explicitly
        gc.collect()
        
        return output
    
    def _get_spectra(self, reader, scan_id):
        ele = reader.get_by_id(scan_id)
        scan_list = ele.get('scanList', {})
        assert scan_list.get('count', None) == 1
        scan = scan_list.get('scan', [])
        assert len(scan) == 1
        scan = scan[0]

        scan_rt = scan.get('scan start time', -1) * 60
        assert scan_rt >= 0

        mz = ele.get('m/z array', np.array([]))
        intensity = ele.get('intensity array', np.array([]))
        return scan_rt, mz, intensity

class ToffeeReader():
    def __init__(
        self,
        tof_fname,
        debug=False,
    ):
        tof_ext = os.path.splitext(tof_fname)[1].lower()
        if tof_ext != '.tof':
            raise ValueError(f'Invalid input file extension (should be .tof): {tof_ext}')
        if not os.path.isfile(tof_fname):
            raise ValueError(f'Input file does not exist: {tof_fname}')
        self.tof_fname = tof_fname

        logger_kwargs = {}
        if debug:
            logger_kwargs['level'] = logging.DEBUG
        self.logger = set_stream_logger(**logger_kwargs)
        self.debug = debug

        self.swath_run = toffee.SwathRun(tof_fname)
        self._ms1_window = None
        self._ms1_spectrum_access = None
        self._ms1_in_memory_spectrum_access = None
        self._ms2_window = None
        self._ms2_spectrum_access = None
        self._ms2_in_memory_spectrum_access = None
        
    def load_ms1_window(self):
        self._ms1_window = self.swath_run.loadSwathMap(toffee.ToffeeWriter.MS1_NAME)
        
    def load_ms1_spectrum_access(self):
        self._ms1_spectrum_access = self.swath_run.loadSwathMapSpectrumAccess(
            toffee.ToffeeWriter.MS1_NAME,
        )
        
    def load_ms1_in_memory_spectrum_access(self):
        self._ms1_in_memory_spectrum_access = self.swath_run.loadSwathMapInMemorySpectrumAccess(
            toffee.ToffeeWriter.MS1_NAME,
        )
        
    def load_ms2_window(self, ms2_name='ms2-050'):
        self._ms2_window = self.swath_run.loadSwathMap(ms2_name)
        
    def load_ms2_spectrum_access(self, ms2_name='ms2-050'):
        self._ms2_spectrum_access = self.swath_run.loadSwathMapSpectrumAccess(ms2_name)
        
    def load_ms2_in_memory_spectrum_access(self, ms2_name='ms2-050'):
        self._ms2_in_memory_spectrum_access = self.swath_run.loadSwathMapInMemorySpectrumAccess(ms2_name)
        
    def extract_spectra(
        self,
        use_ms1,
        in_memory,
        n=1000,
        random_seed=0,
    ):
        np.random.seed(random_seed)
        if use_ms1:
            if in_memory:
                window = self._ms1_in_memory_spectrum_access
                func_name = 'load_ms1_in_memory_spectrum_access'
            else:
                window = self._ms1_spectrum_access
                func_name = 'load_ms1_spectrum_access'
        else:
            if in_memory:
                window = self._ms2_in_memory_spectrum_access
                func_name = 'load_ms2_in_memory_spectrum_access'
            else:
                window = self._ms2_spectrum_access
                func_name = 'load_ms2_spectrum_access'
        if window is None:
            raise RuntimeError(f'Ensure you have run self.{func_name}()')
            
        n_spectra = window.numberOfSpectra()
        spectra_indices = np.random.randint(low=0, high=n_spectra - 1, size=(n,))
        output = []
        for idx in spectra_indices:
            raw_data = window.spectrumByIndex(idx)
            output.append((raw_data.massOverCharge, raw_data.intensities))
        return output
        
    def extract_xic(
        self,
        precursor_mz,
        product_mz_list,
        use_ms1=True,
        ppm_extraction_width=50.0,
        lower_window_overlap=0.0,
        upper_window_overlap=1.0,
    ):
        if use_ms1 and self._ms1_window is None:
            raise RuntimeError('Ensure you have run self.load_ms1_window()')
        if self._ms2_window is None:
            raise RuntimeError('Ensure you have run self.load_ms2_window()')
            
        xic = []
        if use_ms1:
            self.logger.debug('Extract precursor XIC from MS1 window')
            xic.extend(self._extract_xic(self._ms1_window, [precursor_mz], ppm_extraction_width))
            
        self.logger.debug('Extract product XICs from the MS2 window')
        out_lower = self._ms2_window.precursorLowerMz() + lower_window_overlap > precursor_mz
        out_upper = self._ms2_window.precursorUpperMz() - upper_window_overlap <= precursor_mz
        if out_lower or out_upper:
            raise RuntimeError(f'Precursor {precursor_mz} not in MS2 window {self._ms2_window.name()}')
        xic.extend(self._extract_xic(self._ms2_window, product_mz_list, ppm_extraction_width))

        return xic

    def _extract_xic(self, window, product_mz_list, ppm_extraction_width):
        output = []
        for mz in product_mz_list:    
            mz_range = toffee.MassOverChargeRangeWithPPMFullWidth(mz, ppm_extraction_width)
            raw_data = window.extractedIonChromatogram(mz_range)
            output.append((raw_data.retentionTime, raw_data.intensities.sum(axis=1).astype(np.float)))
        # force the garbage collector to manage memory more explicitly
        gc.collect()
        return output

def plot_xics(reader, mz_to_extract):
    xics_to_plot = reader.extract_xic(*mz_to_extract[0], use_ms1=True)
    fig, (ax1, ax2) = plt.subplots(nrows=2, figsize=(15, 5), sharex=True)
    ax1.plot(xics_to_plot[0][0], xics_to_plot[0][1], '-')
    for rt, intensity in xics_to_plot[1:]:
        ax2.plot(rt, intensity, '-')
    ax1.set_ylabel('MS1 XIC Intensity')
    ax2.set_ylabel('MS2 XIC Intensity')
    ax2.set_xlabel('Retention Time (s)')
    fig.tight_layout()
    plt.show()

def plot_spectra(reader, in_memory=None):
    fig, (ax1, ax2) = plt.subplots(nrows=2, figsize=(15, 5), sharex=True)
    for use_ms1, ax in zip([True, False], (ax1, ax2)):
        kwargs = {
            'use_ms1': use_ms1,
            'n': 5,
        }
        if in_memory is not None:
            kwargs['in_memory'] = in_memory
        spectra = reader.extract_spectra(**kwargs)
        for (mz, intensity) in spectra:
#             ax.stem(mz, intensity)
            ax.plot(mz, intensity, 'o', ms=1.0)
    ax1.set_ylabel('MS1 Intensity')
    ax2.set_ylabel('MS2 Intensity')
    ax2.set_xlabel('Mass over charge (Da)')
    fig.tight_layout()
    plt.show()

# %% [code]
mzml_reader = MzMLReader(mzml_fname=f'{base_dir}/ProCan90/mzML/ProCan90-M03-01.mzML')
tof_reader = ToffeeReader(tof_fname=f'{base_dir}/ProCan90/tof/ProCan90-M03-01.tof')

# %% [markdown]
## Initialise the classes

# %% [markdown]
#### `mzML`

To use the `mzML` reader, we need to match the indices in the file to their respective windows.

# %% [code]
%%time

mzml_reader.setup_windows()

# %% [markdown]
#### `toffee`

For the `toffee` reader, we need to load the `SwathMap` classes. Note, as we have filtered the spectral library to only include peptides from window `ms2-050` we only need to load that one. This access pattern is relatively specific to this kind of timing/profiling. For example, in production, there is no need to create both an in-memory and a disk-cached `SpectrumAccess` object.

# %% [code]
%%time

tof_reader.load_ms1_window()

# %% [code]
%%time

tof_reader.load_ms1_in_memory_spectrum_access()

# %% [code]
%%time

tof_reader.load_ms1_spectrum_access()

# %% [code]
%%time

tof_reader.load_ms2_window()

# %% [code]
%%time

tof_reader.load_ms2_in_memory_spectrum_access()

# %% [code]
%%time

tof_reader.load_ms2_spectrum_access()

# %% [markdown]
### Extract Spectrum data

# %% [markdown]
#### `mzML`

# %% [code]
%%time

spectra = mzml_reader.extract_spectra(use_ms1=True)

# %% [code]
%%time

spectra = mzml_reader.extract_spectra(use_ms1=False)

# %% [code]
plot_spectra(mzml_reader)

# %% [markdown]
#### `toffee` -- in memory

# %% [code]
%%time

spectra = tof_reader.extract_spectra(use_ms1=True, in_memory=True)

# %% [code]
%%time

spectra = tof_reader.extract_spectra(use_ms1=False, in_memory=True)

# %% [code]
plot_spectra(tof_reader, in_memory=True)

# %% [markdown]
#### `toffee` -- disk cache

# %% [code]
%%time

spectra = tof_reader.extract_spectra(use_ms1=True, in_memory=False)

# %% [code]
%%time

spectra = tof_reader.extract_spectra(use_ms1=False, in_memory=False)

# %% [code]
plot_spectra(tof_reader, in_memory=False)

# %% [markdown]
### Extract XIC data

# %% [markdown]
#### `mzML`

# %% [code]
%%time

for precursor_mz, product_mz_list in mz_to_extract:
    mzml_reader.extract_xic(precursor_mz, product_mz_list)

# %% [code]
plot_xics(mzml_reader, mz_to_extract)

# %% [markdown]
#### `toffee`

# %% [code]
%%time

for precursor_mz, product_mz_list in mz_to_extract:
    tof_reader.extract_xic(precursor_mz, product_mz_list)

# %% [code]
plot_xics(tof_reader, mz_to_extract)

# %% [code]
```
