## Supplementary material for "Toffee – a highly efficient, lossless file format for DIA-MS": Jupyter notebooks: raptor-part0-transfer_from_resnet50.ipynb

```
# %% [markdown]
# Proof of Concept: Neural Network for detecting peptides

# %% [markdown]
## Introduction

Object detection is the process of identifying and localizing objects in an image. A typical object detection solution takes in an image as input and provides a bounding box on the image where an object of interest is, along with identifying what object the box encapsulates.

This notebook is a proof of concept developed to show that by accessing toffee data in a peptide-centric manner and creating an image from that data, then modern deep learning can be used to detect the retention time of the peptide. 

The notebook was developed for the Amazon SageMaker Object Detection infrastructure to demonstrate how to train an object detection model using the Single Shot multibox Detector ([SSD](https://arxiv.org/abs/1512.02325)) algorithm. It is modified from the [Amazon example](https://github.com/awslabs/amazon-sagemaker-examples/blob/master/introduction_to_amazon_algorithms/object_detection_pascalvoc_coco/object_detection_recordio_format.ipynb).

# %% [markdown]
## Setup

To train the Object Detection algorithm on Amazon SageMaker, we need to setup and authenticate the use of AWS services. To begin with we need an AWS account role with SageMaker access. This role is used to give SageMaker access to your data in S3 will automatically be obtained from the role used to start the notebook.

# %% [code]
%matplotlib inline

# %% [code]
%%time
import boto3
import json
import os

import pandas as pd
import matplotlib.pyplot as plt
import matplotlib.image as mpimg

import sagemaker
from sagemaker import get_execution_role
from sagemaker.amazon.amazon_estimator import get_image_uri

# %% [code]
%%time
role = get_execution_role()
print(role)

# %% [code]
%%time
sess = sagemaker.Session()
training_image = get_image_uri(sess.boto_region_name, 'object-detection', repo_version="latest")
print(training_image)

# %% [markdown]
We also need the S3 bucket that you want to use for training and to store the tranied model artifacts. In this notebook, we require a custom bucket that exists so as to keep the naming clean. You can end up using a default bucket that SageMaker comes with as well.

# %% [code]
bucket = 'procan.users.us-east-1.btully'
prefix = 'Raptor'
run_type = 'zoom'

# %% [markdown]
### Setup S3
We do this using multiple channels. Channels are simply directories in the bucket that differentiate between training and validation data. Let us simply call these directories `train` and `validation`. We will therefore require four channels: two for the data and two for annotations, the annotations ones named with the suffixes `_annotation`.

# %% [markdown]
Set up the data channels -- optionally including a previously trained model. Here, we have previously extracted the data from the toffee files and collected them into an MXNet record file.

# %% [code]
def get_s3_input(name):
    return sagemaker.session.s3_input(
        name,
        distribution='FullyReplicated',
        content_type='application/x-recordio',
        s3_data_type='S3Prefix',
    )

base_dir = 'RaptorTrainingImages/toffee_ml_images'
base_rec_path = f's3://{bucket}/{base_dir}/toffee_ml_images-zoom-easy-only_target-'
data_channels = {
    'train': get_s3_input(base_rec_path + 'train.rec'),
    'validation': get_s3_input(base_rec_path + 'val.rec'),
}

# %% [markdown]
## Training
Now that we are done with all the setup that is needed, we are ready to train our object detector. To begin, let us create a ``sageMaker.estimator.Estimator`` object. This estimator will launch the training job.

# %% [code]
od_model = sagemaker.estimator.Estimator(
    training_image,
    role, 
    train_instance_count=1, 
    train_instance_type='ml.p3.2xlarge',
    train_volume_size=50,
    train_max_run=360000,
    input_mode='File',
    output_path=f's3://{bucket}/{prefix}/{run_type}/part0',
    sagemaker_session=sess,
)

# %% [markdown]
The object detection algorithm at its core is the [Single-Shot Multi-Box detection algorithm (SSD)](https://arxiv.org/abs/1512.02325). This algorithm uses a `base_network`, which is typically a [VGG](https://arxiv.org/abs/1409.1556) or a [ResNet](https://arxiv.org/abs/1512.03385). For this proof of concept we have taken a pre-trained ResNet-50 model. It also has a lot of options for hyperparameters that help configure the training job. The next step in our training, is to setup these hyperparameters and data channels for training the model. Consider the following example definition of hyperparameters. See the SageMaker Object Detection [documentation](https://docs.aws.amazon.com/sagemaker/latest/dg/object-detection.html) for more details on the hyperparameters.

One of the hyperparameters here for instance is the `epochs`. This defines how many passes of the dataset we iterate over and determines that training time of the algorithm, we have selected `60` epochs so that training should take less than a day.

# %% [code]
s3_client = boto3.client('s3', region_name='us-east-1')
s3_transfer = boto3.s3.transfer.S3Transfer(client=s3_client)

fname = 'toffee_ml_images-zoom-easy-only_target-train.lst'
s3_transfer.download_file(
    bucket,
    f'{base_dir}/{fname}',
    fname,
)
lst = pd.read_csv(fname, sep='\t', names='Idx,HeaderSize,HeaderShape,Class,Xmin,Ymin,Xmax,Ymax,FileName'.split(','))

num_training_samples = lst.shape[0]
f'num_validation_samples = {num_training_samples}'

# %% [code]
od_model.set_hyperparameters(
    base_network='resnet-50',
    use_pretrained_model=1,
    num_classes=1,
    mini_batch_size=32,
    epochs=60,
    learning_rate=0.001,
    lr_scheduler_step='3,6',
    lr_scheduler_factor=0.1,
    optimizer='sgd',
    momentum=0.9,
    weight_decay=0.0005,
    overlap_threshold=0.75,
    nms_threshold=0.45,
    image_shape=300,
    label_width=300,
    num_training_samples=num_training_samples,
)

# %% [markdown]
We have our `Estimator` object, we have set the hyperparameters for this object and we have our data channels linked with the algorithm. The only remaining thing to do is to train the algorithm. The following cell will train the algorithm. Training the algorithm involves a few steps. Firstly, the instances that we requested while creating the `Estimator` classes are provisioned and are setup with the appropriate libraries. Then, the data from our channels are downloaded into the instance. Once this is done, the training job begins. The provisioning and data downloading will take time, depending on the size of the data. Therefore it might be a few minutes before we start getting data logs for our training jobs. The data logs will also print out Mean Average Precision (mAP) on the validation data, among other losses, for every run of the dataset once or one epoch. This metric is a proxy for the quality of the algorithm. 

Once the job has finished a "Job complete" message will be printed. The trained model can be found in the S3 bucket that was setup as `output_path` in the estimator.

# %% [code]
od_model.fit(inputs=data_channels, logs=False)

# %% [markdown]
## Deploy

Once the training is done, we can deploy the trained model as an Amazon SageMaker real-time hosted endpoint. This will allow us to make predictions (or inference) from the model. Note that we don't have to host on the same insantance (or type of instance) that we used to train. Training is a prolonged and compute heavy job that require a different of compute and memory requirements that hosting typically do not. We can choose any type of instance we want to host the model. In our case we chose the ml.p3.2xlarge instance to train, but we choose to host the model on the less expensive cpu instance, ml.m4.xlarge. The endpoint deployment can be accomplished as follows:

# %% [code]
object_detector = od_model.deploy(
    initial_instance_count=1,
    instance_type = 'ml.m4.xlarge',
)
object_detector.content_type = 'image/png'

# %% [markdown]
## Inference

Now that the trained model is deployed at an endpoint that is up-and-running, we can use this endpoint for inference.

We can see in these results that there are a significant number of false negatives. However, with minimal effort and adjustment to the model architecture, the true positives show that there is a high likelihood that deep-learning will prove an effective method for peptide detection from toffee data.

# %% [code]
def visualize_detection(img_file, dets, thresh=0.6):
        """
        visualize detections in one image
        Parameters:
        ----------
        img : numpy.array
            image, in bgr format
        dets : numpy.array
            ssd detections, numpy.array([[id, score, x1, y1, x2, y2]...])
            each row is one object
        classes : tuple or list of str
            class names
        thresh : float
            score threshold
        """
        f, ax = plt.subplots(num=0)
        img = mpimg.imread(img_file)
        ax.imshow(img)
        height = img.shape[0]
        width = img.shape[1]
        for det in dets:
            (klass, score, x0, y0, x1, y1) = det
            if score < thresh:
                continue
            cls_id = int(klass)
            xmin = int(x0 * width)
            ymin = int(y0 * height)
            xmax = int(x1 * width)
            ymax = int(y1 * height)
            rect = plt.Rectangle(
                (xmin, ymin),
                xmax - xmin,
                ymax - ymin,
                fill=False,
                edgecolor='maroon',
                linewidth=2,
            )
            ax.add_patch(rect)
            ax.text(
                xmin,
                ymin - 2,
                f'PQM {score:.3f}',
                bbox=dict(
                    facecolor='maroon',
                    alpha=0.5
                ),
                fontsize=12,
                color='white',
            )
        f.savefig(img_file.replace('.png', '-prediction.png'))
        plt.show()

# %% [code]
val_fname = 'toffee_ml_images-zoom-easy-only_target-val.lst'
# val_fname = 'toffee_ml_images-zoom-only_target-val.lst'
s3_transfer.download_file(
    bucket,
    f'{base_dir}/{val_fname}',
    val_fname,
)
val_lst = pd.read_csv(val_fname, sep='\t', names='Idx,HeaderSize,HeaderShape,Class,Xmin,Ymin,Xmax,Ymax,FileName'.split(','))
val_lst.head()

# %% [code]
import os

threshold = 0.2
num_printed = 0

for i, row in val_lst.sample(frac=1, random_state=0).iterrows():
    for decoy in [False, True]:
        fname = row.FileName
        if decoy:
            fname = fname.replace('.png', '_DECOY.png')
        awsdir = os.path.dirname(fname)
        fname = os.path.basename(fname)
        if not os.path.exists(fname):
            try:
                s3_transfer.download_file(
                    bucket,
                    f'RaptorTrainingImages/{awsdir}/{fname}',
                    fname,
                )
            except Exception:
                break
         
        if not decoy:
            print('-' * 80, flush=True)
            print(row.FileName)
            print('\tTARGET -- there should be a detection', flush=True)
        else:
            print('\tDECOY -- there should be NO detection', flush=True)

        with open(fname, 'rb') as image:
            f = image.read()
            b = bytearray(f)

        detections = json.loads(object_detector.predict(b))
        visualize_detection(fname, detections['prediction'], thresh=threshold)

        os.remove(fname)

    num_printed += 1
    if num_printed == 10:
        break

# %% [markdown]
## Delete the Endpoint

Having an endpoint running will incur some costs. Therefore as a clean-up job, we should delete the endpoint.

# %% [code]
sess.delete_endpoint(object_detector.endpoint)
```
